# HSF1 controls transcriptional programs that establish thalamostriatal shaft synaptic architecture and preserve cognitive flexibility

**DOI:** 10.64898/2026.08.03.742053

**Authors:** Nicholas B Rozema, Nicole Zarate, Rachel Mansky, Persephone Gu, Kinsey Gerlach, Aishwarya Bhowmik, James H Cho, Ying Zhang, Arif Hamid, Steven M Graves, Rocio Gomez-Pastor

## Abstract

Cognitive flexibility (CF) declines during aging and is further impaired in neurodegenerative diseases such as Huntington’s disease (HD), yet the molecular mechanisms underlying these deficits remain poorly understood. Thalamostriatal (T-S) synapses are critical for CF, and we previously identified Heat Shock Factor 1 (HSF1) as a regulator of T-S density in HD. However, how HSF1 regulates T-S synapses and whether it modulates cognitive flexibility (CF) remained unclear. Here, we combined HSF1 ChIP-seq, transcriptomics, synapto-proteomics, targeted genetic manipulations and behavioral analyses to study how HSF1 regulates T-S synapses and CF. We found HSF1 directly controls a transcriptional program governing postsynaptic architecture and actin cytoskeletal dynamics, which are disrupted in aging and HD. Loss of HSF1 drives selective destabilization of actin patches at T-S shaft synapses and impaired CF decline. Our results underscore a novel function for HSF1 in the regulation of striatal neural circuits with essential implications in the neurobiology of cognitive flexibility.

## Introduction

Cognitive flexibility (CF) is a core aspect of executive functioning that consists of an individual’s ability to adapt their behavior and thinking in response to changes in the environment. Impairments in CF occur as a normal result of cognitive aging^1–5^ and lead to difficulty in functional independence, communication, and socialization in the elderly. Across numerous neurodegenerative diseases, including Huntington’s disease (HD), the age-related decline in CF is accelerated and exacerbated^1,6–12^. However, the molecular mechanisms responsible for CF decline in both aging and neurodegeneration are not well understood. A majority of studies related to CF have focused on the hippocampus and cortex due to their established roles in learning and memory^13–15^. However, changes within glutamatergic signaling in the dorsal striatum, a key brain area involved in motor control and the primary site of neurodegeneration in HD, have long been implicated in regulating CF in both elderly individuals and HD patients^16–26^.

The striatum’s function is regulated by two major excitatory glutamatergic inputs: corticostriatal (C-S) and thalamostriatal (T-S). Medium spiny neurons (MSN) in the striatum rely equally on these excitatory inputs^27–29^, although T-S and C-S synapses present distinct electrophysiological properties and therefore influence MSNs differently^30,31^. Importantly, prior research has shown that T-S synapses play a key role in modulating behavioral flexibility, including CF^23,32–36^. Multiple studies have reported that T-S synapses are among the earliest synaptic populations affected in HD, degenerating before C-S synapses and prior to the onset of overt motor symptoms^37–40^. However, whether T-S synapse loss contributes to the decline in CF observed in HD remains unresolved^6^. In contrast, although age-related impairments in CF are well documented, it is currently unknown whether T-S synapses are similarly vulnerable during normal aging, as T-S synapse density in the aging striatum has not yet been systematically investigated. Consequently, the potential contribution of T-S synapse degeneration to CF decline in the aging striatum remains unexplored. These critical knowledge gaps highlight the need for a deeper understanding of the mechanisms underlying T-S synapse vulnerability and the extent to which T-S degeneration contributes to cognitive dysfunction in both aging and HD.

Heat Shock Factor 1 (HSF1) is a stress protective transcription factor classically associated with the heat shock response (HSR) through the regulation of molecular chaperones^41,42^. Accumulating evidence has revealed that HSF1 is able to regulate a broad network of additional genes depending on the context. These include genes essential to cell cycle regulation, glucose metabolism, inflammatory responses, cell wall and cytoskeletal stability, metabolism, and the development and maintenance of neuronal, reproductive, and sensory organs^43–48^. While emerging evidence highlights the importance of HSF1 in the regulation of neuronal and synaptic fate, particularly in the context of neurodegenerative diseases^42,48–61^, its role has traditionally been viewed in a rather simplistic manner and primarily attributed to the control of chaperones and other protein quality control components. This perspective has been influenced by the limited number of studies examining the broader regulatory functions and diverse physiological roles of HSF1 under non stressful conditions and by the scarcity of studies involving direct manipulation of HSF1 in the brain.

Previous studies have shown that HSF1 directly regulates the expression of some synaptic genes (PSD95, SAP97 and BDNF)^62–64^, thereby establishing a connection between HSF1 and the maintenance of synaptic stability through direct regulation of structural synaptic components and neurotrophic factors^62^. Additional studies have further supported the importance of HSF1 in synapse biology. Consistent with this function, HSF1-deficient mice exhibit impaired hippocampal synaptogenesis^65^. Moreover, HSF1 haploinsufficient mice display an age-dependent and selective loss of T-S synapses^62^, highlighting a critical role for HSF1 in synaptic specialization and maintenance within the basal ganglia. Importantly, HSF1 levels are reduced in both the aging striatum and HD, and experiments aimed at restoring the levels of HSF1 in HD mice prevented T-S synapse loss^49,62^. However, because HSF1 also plays important roles during development^66,67^, and prior studies have relied largely on systemic HSF1 deletion models, the specific contribution of HSF1 to the maintenance and specialization of synapses in the mature brain remains unresolved. As a result, it is still unknown whether HSF1 directly regulates the stability of T-S synapses in adulthood and, consequently, influences age-related CF decline.

In this study, we sought to define the relationship between HSF1, T-S synapse stabilization, and the regulation of CF. Using an integrated approach that combined ChIP-seq, RNA-seq, synapto-proteomics, behavioral analyses, and brain-specific genetic manipulations of HSF1, we uncovered a previously unrecognized role for HSF1 in the maintenance of axodendritic (shaft) synaptic connections within the T-S pathway. Mechanistically, HSF1 regulated the expression of actin-binding proteins and cytoskeletal regulators that control the formation of dendritic actin patches required for T-S shaft synapse development and stability. Importantly, brain-specific reductions in HSF1 selectively impaired T-S shaft synapses and were accompanied by deficits in CF. Collectively, these findings establish HSF1 as a critical regulator of striatal synaptic architecture and identify a novel molecular pathway linking synaptic stability to cognitive function with important implications for understanding CF decline during aging and neurodegenerative disease.

## Results

### Thalamostriatal synapses are selectively lost in conditions of *Hsf1* dysfunction and deficiency

Striatal synaptic dysfunction and synaptic loss are hallmarks of both normative aging and HD pathology^68–72^, but whether there are common mechanisms responsible for similar synaptic alterations in these different conditions is unknown. These conditions also have in common a critical decline in the levels of HSF1 in the striatum^49,62^. Our lab has previously shown that dysfunction and degradation of HSF1 in HD is related to pathway-specific T-S synapse loss since restoring the levels of HSF1 in the HD mouse model zQ175 increased T-S synapse density^49^. We also demonstrated that HSF1 protein levels decline in the dorsal striata of WT mice as a function of normative aging^62^. Therefore, we first investigated whether WT mice show age-dependent and circuit-specific changes in total synaptic density in the dorsal striatum. Utilizing confocal microscopy, we found an age-dependent increase in the total density of C-S synapses in the dorsal striatum, which are mostly axospinous^73^, measured by colocalization of the presynaptic marker VGLUT1 and postsynaptic marker PSD95 (**Figure 1A-C**). This was accompanied by a significant decrease in the total density of T-S synapses (colocalization of VGLUT2/PSD95) represented by both axospinous and shaft synapses (**Figure 1D-F**). As previously shown^49,62^, these time points aligned with a significant decrease in total HSF1 levels (**Supplementary Figure 1A,B**). Consistent with previous data^37,38,49,62^, we confirmed an early and selective loss of T-S synapses in zQ175 mice (hereafter, HD mice) using two independent postsynaptic markers (PSD95 and Homer1) (**Figure 1G-J, Supplementary Figure 1C-E**). Notably, this initial synapse loss in HD mice occurs before detectable reductions in HSF1 protein levels (**Supplementary Figure 1A,B)**. However, we previously demonstrated that HSF1 is functionally inactivated prior to its degradation^49^ suggesting that impaired HSF1 activity, rather than loss of protein abundance, coincides with the onset of T-S synapse degeneration.

**Figure 1.**
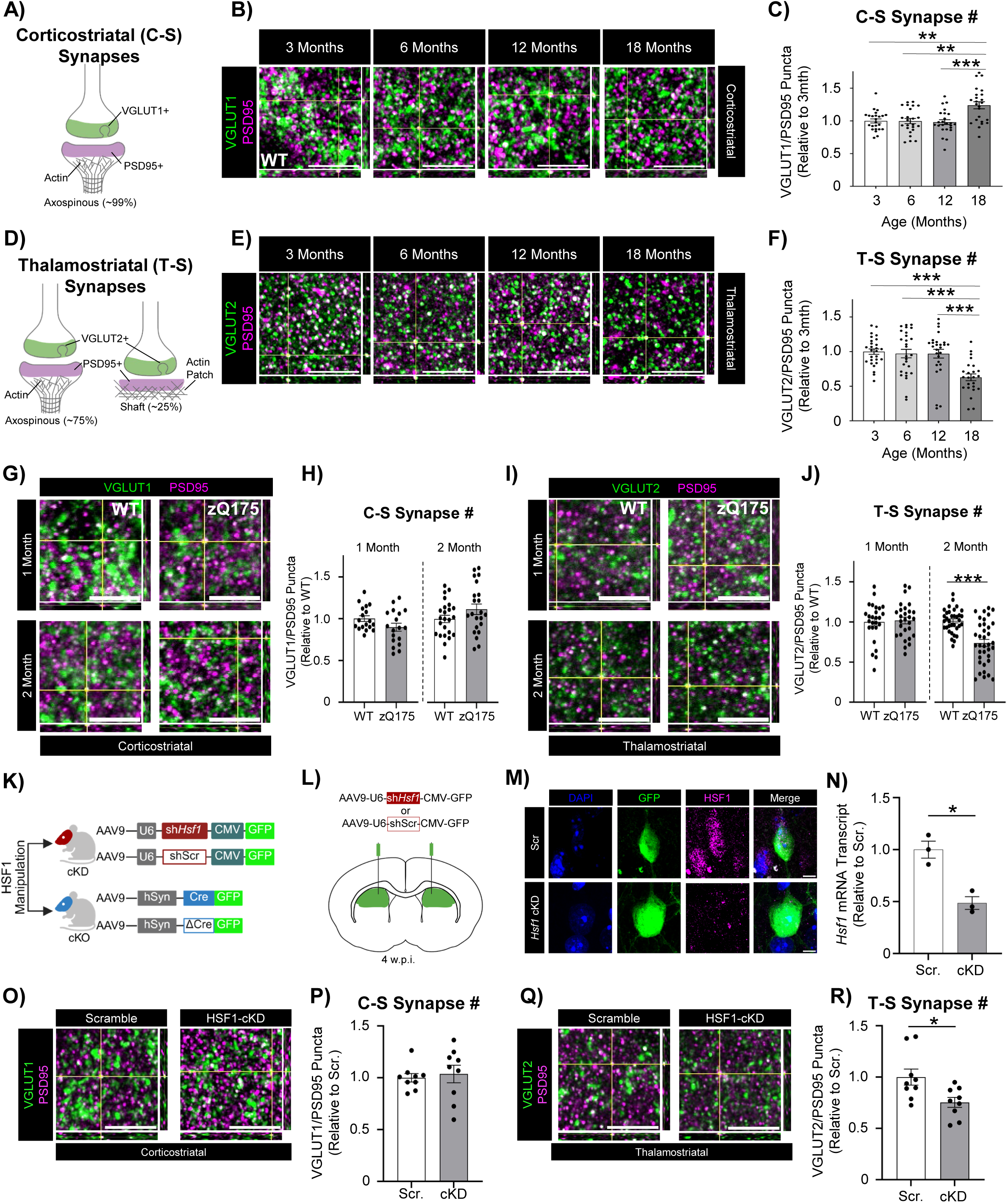
Selective loss of thalamostriatal synapses occurs in conditions of HSF1 deficiency. **(A)** Schematic of axospinous (spiny) C-S synapses labeled by presynaptic VGLUT1. **(B)** Representative images of C-S synapses identified by colocalization of VGLUT1/PSD95 in 3-month-, 6-month-, 12-month-, and 18-month-old wildtype striatum. **(C)** Quantification of C-S synapses from B (n=9 mice/3 slices/group). **(D)** Schematic of axospinous (spiny) and shaft T-S synapses labeled by presynaptic VGLUT2. **(E)** Representative images of T-S synapses identified by colocalization of VGLUT2/PSD95 in 3-month, 6-month, 12-month, and 18-month-old wildtype striatum. **(F)** Quantification of T-S synapses from E (n=9 mice/3 slices/group). **(G)** Representative images of C-S synapses from 1-month and 2-month-old wildtype and HD striatum. **(H)** Quantification of C-S synapses from G (n=6 mice/3 slices/group). **(I)** Representative images of T-S synapses from 1-month and 2-month-old wildtype and HD striatum. **(J)** Quantification of C-S synapses from I (n=8 mice/3 slices/group). **(K)** HSF1 manipulation methods used in adult 6-month-old mice via AAV intracranial injection for HSF1-cKD and HSF1-cKO. **(L)** Intracranial injection scheme used for HSF1-cKD. **(M)** Representative images of HSF1-cKD in striatal neurons. **(N)** *Hsf1* mRNA transcript levels from bulk striatal extracts of control (Scr) and HSF1-cKD mice (sh*Hsf1*) (n=3 mice/group). **(O)** Representative images of C-S synapses from HSF1-cKD striatum. **(P)** Quantification of C-S synapses from O. **(Q)** Representative images of T-S synapses from HSF1-cKD striatum (n=3 mice/3 slices/group). **(R)** Quantification of T-S synapses from Q (n=3 mice/3 slices/group). Orthogonal views are provided for each representative synapse image as confirmation of colocalization. Data points in synapse density analysis represent a single slice. Scale Bars: 5 μm. Error bars ±SEM. Ordinary one-way ANOVA (C,F). Two-Tailed Student’s t-test (H,J,P,R). * p<0.05, ** p<0.005, *** p<0.0005.

Previous work from our laboratory using *Hsf1* haploinsufficient (+/-) mice demonstrated that reduced HSF1 levels, and therefore reduced activity, are sufficient to decrease T-S synapse density in healthy mice^62^. However, because HSF1 plays essential roles during development, these studies could not establish a direct role for HSF1 in maintaining the stability of T-S synapses in the adult brain. Instead, the observed synaptic alterations may have arisen, at least in part, from neurodevelopmental defects resulting from the constitutive systemic reduction of HSF1. To resolve this issue, we utilized a viral-mediated conditional knockdown approach (HSF1-cKD) to selectively deplete HSF1 in the fully developed adult brain. We used an AAV9 expressing a short-hairpin RNA targeted against *Hsf1* (AAV9-U6-sh*Hsf1*-CMV-GFP) or a scrambled RNA (AAV9-U6-shScr-CMV-GFP) as control injected bilaterally into the dorsal striatum of 6-month-old WT mice (**Figure 1K-N; Supplementary Figure 1F,G**). We found that HSF1-cKD does not significantly impact the density of C-S synapses (**Figure 1O-P**), but significantly depleted the density of T-S synapses (**Figure 1Q-R**). We confirmed this finding with a conditional HSF1 KO mouse (HSF1-cKO). 6-month-old *Hsf1*^fl/fl^ mice injected with an AAV9 expressing Cre under the human Synapsin promoter (AAV9-hSyn-Cre-GFP) also showed a selective decrease of T-S synapses, but not C-S synapses compared to control *Hsf1*^fl/fl^ mice injected with the catalytic dead ΔCre (AAV9-hSyn-ΔCre-GFP) (**Figure 1K**, **Supplementary Figure 1H-N**). All together, these data demonstrate that HSF1 depletion in the adult brain is sufficient to drive a selective destabilization of T-S synapses in the dorsal striatum.

### HSF1 preferentially occupies synaptic gene networks in the striatum that are disrupted in HD

Given the apparent relationship between striatal HSF1 and the regulation of T-S synapse density, we next sought to examine whether HSF1 may engage in direct regulatory control of genes important for synaptic stabilization and function. We first sought to characterize the HSF1 DNA-binding landscape in the striatum to identify genes that are directly regulated by HSF1 and may contribute to the maintenance of T-S synapses. HSF1 DNA-binding profiles have been established in cancer cells and other conditions^74–76^ and show great versatility and context-dependence related to stress response, metabolism, and angiogenesis, but have never been explored in the brain. To capture potential changes in HSF1 target gene occupancy during critical time points of striatal development and maturation, we conducted HSF1 chromatin immunoprecipitation (ChIP)-sequencing during postnatal development (1-month) and after maturation (2-months) (**Figure 2A**). These points were also selected because they span the period preceding and coinciding with the early loss of T-S synapses in HD mice, enabling us to identify HSF1-dependent transcriptional programs that may underline synaptic vulnerability in both the mature brain and HD.

**Figure 2.**
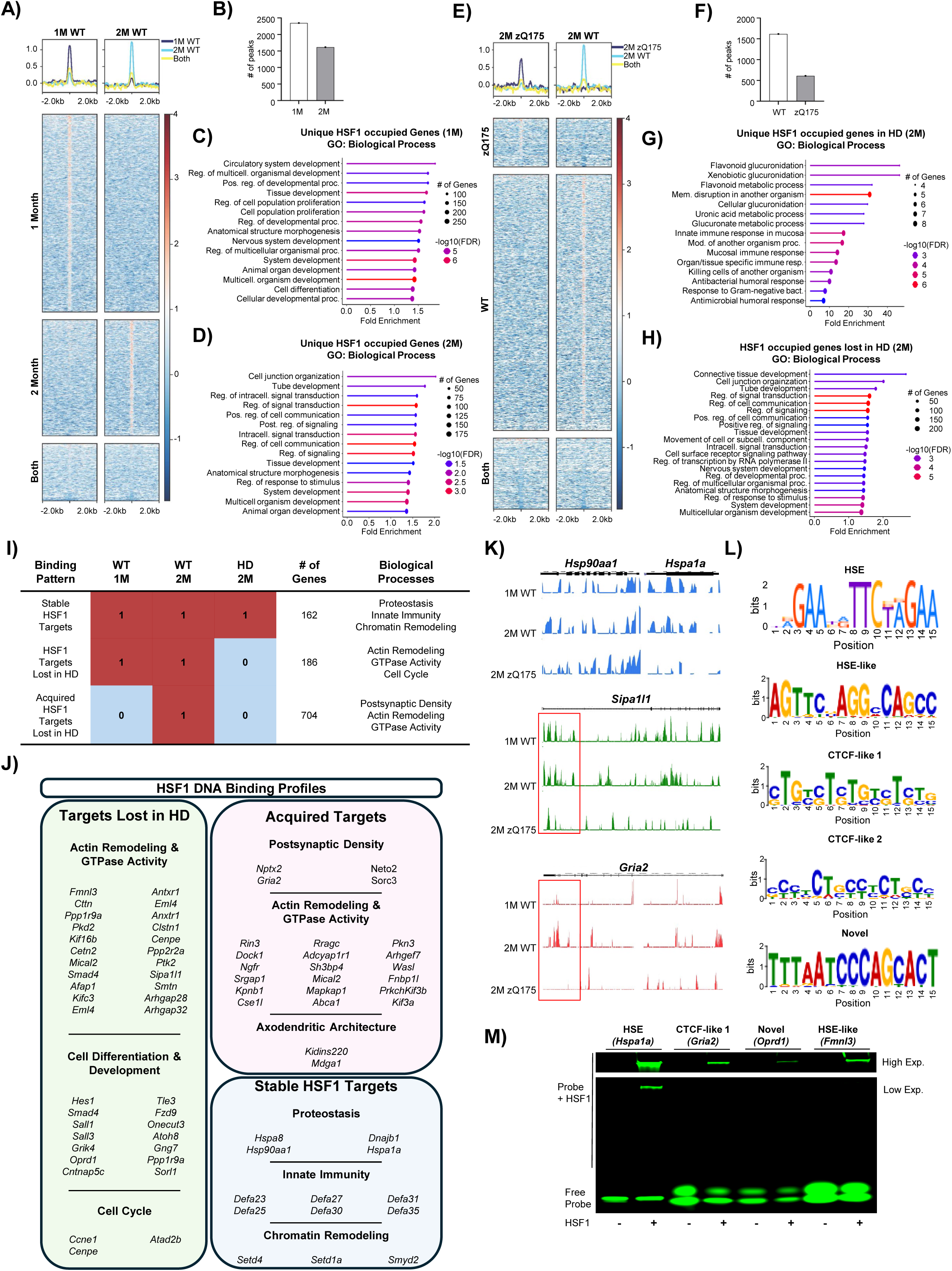
HSF1 DNA occupancy is developmentally remodeled toward synaptic genes and fails to mature in Huntington’s disease. **(A)** HSF1 ChIP-seq profile plots and heatmaps from 1-month and 2-month-old WT mouse striatum. **(B)** Total peaks detected in 1-month and 2-month-old HSF1 ChIP-seq datasets from WT mice. **(C)** Gene ontology for cellular component of genes detected in 1-month-old WT HSF1 ChIP-seq dataset. **(D)** Gene ontology for cellular component of genes detected in 2-month-old WT HSF1 ChIP-seq dataset. **(E)** HSF1 ChIP-seq profile plots and heatmaps from 2-month-old WT and HD mouse striatum. **(F)** Total peaks detected in 2-month-old WT and HD ChIP-seq datasets. **(G)** Gene ontology for biological process of genes detected in 2-month-old HD HSF1 ChIP-seq dataset. **(H)** Gene ontology for biological process of genes detected in 2-month-old WT HSF1 ChIP-seq dataset not bound in HD mice. **(I)** Distinct HSF1 DNA binding profiles detected across 1-month- and 2-month-old WT striatum and 2-month-old HD mice. **(J)** Representative genes found in each unique HSF1 DNA binding profile. **(K)** Examples of HSF1 binding to the genes *Hsp90aa1* and *Hspa1a* (Stable HSF1 Targets), *Sipa1l1* (HSF1 Target Lost in HD), and *Gria2* (Developmentally Acquired HSF1 Target Lost in HD). Red boxes indicate areas of particularly robust HSF1 binding differences. **(L)** Motifs identified through *de novo* motif enrichment across all ChIP-seq datasets. **(M)** HSF1 EMSA for binding sites of *Gria2* (CTCF-like 1), *Oprd1* (Novel), *Fmnl3* (HSE-like) and *Hsp70* (HSE control). n=6-8 mice/group.

Surprisingly, HSF1 ChIP-seq in WT mice at 1 vs 2 months of age showed large numbers of bound genes at both timepoints (**Figure 2B**). Specifically, at 1-month we detected 2,347 peaks aligning to 1,429 genes and at 2-months we detected 1,616 peaks aligning to 1,082 genes (**Figure 2B, Supplementary File 1**). The binding patterns we detected were also distinct depending on developmental stage with 1,045 unique genes bound at 1-month, 704 unique genes at 2-months, and only 186 shared genes between the two timepoints (**Supplementary File 1)**. Using the ShinyGO bioinformatics tool^77^, we determined that genes uniquely bound in the 1-month-old striatum were enriched for biological processes involved in tissue development and cell proliferation (**Figure 2C**) consistent with HSF1’s previously described roles in neurodevelopment in other brain regions^65,78,79^. However, at 2-months the binding profile of HSF1 changed drastically and showed enrichment in genes involved in signal transduction and cell junction organization (**Figure 2D**). When conducting gene ontology analyses based on cellular component, we found that although both time points presented synaptic components, the distinction laid in a preferential projection-promoting profile at 1-month during striatal postnatal development and a somatodendritic, glutamatergic, and postsynaptic specialization profile at 2-months (**Supplementary Figure 2A,D**). This data suggests a wide participation of HSF1 in the regulation of key neuronal programs in the striatum that shifts from neuronal development and differentiation towards a specialized glutamatergic synaptic stabilization in the mature striatum.

We next investigated whether the genomic occupancy of HSF1 is altered in the striatum of HD mice, potentially contributing to the selective vulnerability of T-S synapses in HD. To this end, we performed HSF1 ChIP-seq in the striatum of 2-month-old HD mice, an early disease stage at which T-S synapse density is significantly reduced (**Figure 1I,J**) and compared the resulting HSF1 DNA binding profile with that of age-matched WT mice (**Figure 2E**). HSF1 chromatin occupancy was profoundly altered in the HD striatum. We identified a total of 610 HSF1-binding peaks associated with 277 annotated genes in zQ175 mice in contrast to the 1,616 peaks found in WT mice (**Figure 2F**). Among those 610 identified peaks, we found a subset of 49 genes that were uniquely occupied in the HD striatum and not in WT mice and 228 genes that were shared between the two genotypes (**Supplementary File 1**). Genes bound in HD mice corresponded to biological processes related to xenobiotic/detoxification metabolism and innate immune response consistent with a disease-associated redistribution of HSF1 binding toward inflammatory gene programs (**Figure 1G**). These genes did not reveal any significant cellular component. Importantly, we found that 805 genes lost HSF1 occupancy in HD mice (**Figure 2E, F**). Those genes were significantly enriched for pathways related to synaptic function, including cell junction organization and the regulation of synaptic signaling, corresponding to the functional categories prominently represented among HSF1 targets in the WT striatum (**Figure 2H**).

Based on our ChIP-seq analysis we identified 3 distinct HSF1 gene occupancy profiles across postnatal development, maturity, and disease state (**Figure 2I-K**): (i) *Stable HSF1 targets* corresponds to genes that are occupied by HSF1 regardless of development stage or disease state and include genes related to proteostasis and molecular chaperones (*Hsp90aa1, Hspa1a, Hspa5, Dnajb1*), innate immunity (*Defa23/26/27/30/31/35*), and chromatin remodeling (*Setd4, Smyd2, Setd1a*) amongst other processes. (ii) *Stable HSF1 targets lost in HD* correspond to stable HSF1 targets across development stages (1- and 2-months-old WT) that are lost in HD and includes genes involved in cytoskeletal/actin remodeling and GTPase activity (ex. *Sipa1l1, Ppp1r9a, Arhgap28, Arghap32*), cell differentiation (ex. *Hes1, Smad4, Sall1/3*), and cell cycle regulation (*Ccne1, Cenpe, Atdad2b*). (iii) Acquired HSF1 targets lost in HD correspond to genes occupied by HSF1 only in the matured striatum and that are lost in HD and includes genes involved in actin remodeling and GTPase activity (ex. *Dock1, Srgap1, Arhgef7, Wasl, Kif3a/b*), as well as specialized postsynaptic density genes (*Nptx2, Gria2, Neto2, Sorc3, Oprd1*) and genes involved in establishing axodendritic architecture (*Kidins220, Mdga1*). HSF1 occupancy onto a subset of these targets (*Gria2* and *Oprd1*) was validated via HSF1 ChIP in control striatal ST*Hdh*^Q^^7/Q^^7^ and HD ST*Hdh*^Q^^111/Q^^7^ murine cells (**Supplementary Figure 3A-B**). Together, our data indicates that HSF1 is actively engaged in binding to a plethora of genes under physiological, non-stressful conditions in the mammalian central nervous system, first associating with genes involved in broad neurodevelopment and later in synaptic formation and specialization. Importantly, HD pathology appears to redirect HSF1 occupancy away from genes involved in synaptic function toward genes associated with innate immune and stress-response programs. This redistribution coincides with the earliest detectable loss of T-S synapses in HD zQ175 mice, suggesting that disruption of the normal HSF1 synaptic regulatory network may contribute to the selective vulnerability of T-S synapses observed in HD, consistent with the synaptic deficits we observed following HSF1 deficiency.

To more closely examine if HSF1 binding sites detected in our ChIP-Seq dataset correspond to canonical heat shock elements (HSE; GAAnnTTCnnGAA) or if they represent other uncharacterized HSF1 binding motifs, we employed *de novo* motif enrichment analysis^80^. We found five different motifs that could be grouped into HSE and HSE-like motifs, two CTCF-like motifs which align more closely with the previously reported binding site of the CCCTC-binding factor (CTCF)^81^ (consensus motif: CCGCGnGGnGGCAG), and a novel uncharacterized motif (TTTAATCCCAGCACT) (**Figure 2L**). CTCF has been previously described to form protein-protein interactions with HSF1 under non-stressed, stressed, and HD conditions, and concomitant occupancy of HSF1 and CTCF on DNA regions often results in transcriptional repression of the target genes^82^. Interestingly, CTCF-like motifs were more enriched among the HSF1 occupied targets than the HSE/HSE-like motifs (**Supplementary Figure 3C**). However, genomic regions that lost HSF1 occupancy in the HD striatum did not exhibit preference for any specific DNA binding motif, suggesting that the loss of HSF1 binding is not associated with a binding impairment to a particular sequence element (**Supplementary Figure 3D**). To validate direct HSF1 binding to genomic regions identified by ChIP-seq, we performed electrophoretic mobility shift assays (EMSAs) using IR800-labeled double-stranded DNA probes corresponding to representative HSF1-bound sequences. Specifically, we tested *Gria2* (CTCF-like 1 motif), *Oprd1* (Novel), *Fmnl3* (HSE-like motif), and *Hspa1a* (HSE) as a positive control. As expected, HSF1 exhibited strong binding to the *Hspa1a* (Hsp70) HSE probe. Importantly, however, HSF1 also bound each of the three other motifs identified by our ChIP-seq analysis, although these interactions were weaker than *Hspa1a*:HSE (**Figure 2M**). These findings demonstrate that HSF1 is capable of directly recognizing multiple non-canonical motifs, supporting the notion that HSF1 occupies a broader repertoire of genomic binding sites than previously appreciated. The enrichment of CTCF-like motifs among HSF1-bound regions raises the possibility that cooperative interactions with CTCF or other chromatin-associated factors enhance HSF1 recruitment and stabilization at these loci *in vivo*.

### Loss of HSF1 causes abnormal expression of structural synaptic genes

We next sought to confirm that synaptic genes occupied by HSF1 are directly regulated by HSF1. To accomplish this, we performed RNA-sequencing (RNA-Seq) in 2-month-old WT and *Hsf1^+/-^* striata (**Figure 3A**). We chose Hsf1 happloinsufficient mice to better mimic the conditions of chronic depletion seen in the brains of HD and aged mice. We detected 1,626 differentially expressed genes (DEGs) of which 741 transcripts were significantly down-regulated and 885 transcripts were significantly up-regulated in *Hsf1^+/-^* compared to WT (**Figure 3B,C, Supplementary File 2**). Across all DEGs, transcript changes were nearly evenly distributed between upregulated (54%) and downregulated (46%) genes (**Figure 3D**). Gene Ontology (GO) analysis for Cellular Component revealed that upregulated genes were predominantly enriched for ribosomal subunits and protein translation-associated complexes although they also included a synapse category (**Figure 3E, Supplementary Figure 3A-C**). The downregulated genes were enriched for synaptic and neuronal cellular components, especially somatodendritic (**Figure 3F, Supplementary Figure 3D-F**). When the analysis was restricted to only genes annotated with synaptic function based on GO, a clear bias emerged: 64% of synaptic DEGs were upregulated in *Hsf1*^+/-^ vs WT mice while only 36% were downregulated (**Figure 3D**). This enrichment of upregulated synaptic transcripts suggested that HSF1 played a crucial role as a transcriptional repressor of synaptic genes, directly or indirectly.

**Figure 3.**
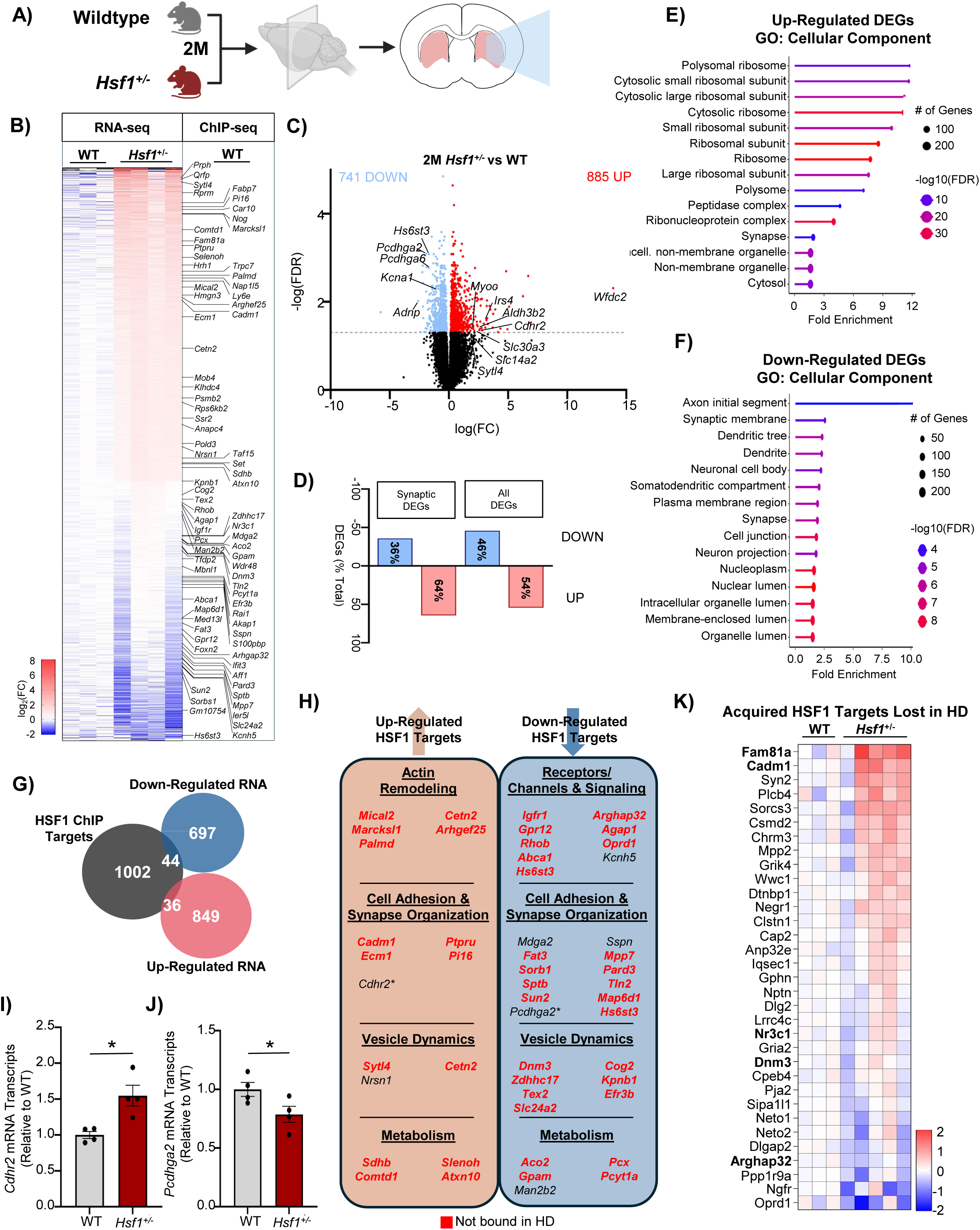
Loss of HSF1 activity results in transcriptional dysregulation of synaptic genes. **(A)** Schematic of RNA-seq experiment in 2-month-old WT and *Hsf1^+/-^* mice. **(B)** Heat map of all differentially expressed genes (DEGs) in 2-month-old *Hsf1^+/-^* mouse striatum annotated with direct HSF1 targets from 2-month-old HSF1 ChIP-seq. **(C)** Volcano plot of DEGs in 2-month-old *Hsf1^+/-^* compared to WT mouse striatum. **(D)** Overview of down- and up-regulation of all DEGs in 2-month-old *Hsf1^+/-^* compared to WT striatum and synaptic DEGs. **(E)** Gene ontology for cellular component of genes that are up-regulated in 2-month-old *Hsf1^+/-^* striatum. **(F)** Gene ontology of cellular component of genes that are down-regulated in 2-month-old *Hsf1^+/-^* striatum. **(G)** Venn diagram of DEGs in 2-month-old *Hsf1^+/-^* mouse striatum and genes detected in 2-month-old WT HSF1 ChIP-seq. **(H)** Categories of DEGs that are direct HSF1 targets in ChIP-seq. Red text indicates transcripts not bound in 2-month-old HD mice ChIP-seq. * indicates non-HSF1 targets validated by RT-qPCR. **(I-J)** qPCR validation of DEGs in *Hsf1^+/-^* mice (*Cdhr2* and *Pcdhga2)*. **(K)** Heat map of direct HSF1 targets associated with postsynaptic density formation and function. DEGs: q<0.05 determined by DESeq2 (v1.46.0). n=3-5 mice/group (RNA-seq); 4 mice/group (RT-qPCR). Two-Tailed Student’s t-test (I,J) *p<0.05. Error bars ±SEM.

When we compared the DEGs obtained by RNA-seq and HSF1 occupied genes identified by ChIP-seq we found a total of 80 transcripts that represent putative direct HSF1 targets and are differentially expressed in *Hsf1*^+/-^ mice (**Figure 3G**). Notably, HSF1-regulated transcripts displayed distinct directional patterns, with increased expression of genes associated with actin remodeling, cell adhesion and synapse organization, vesicle dynamics, and metabolism, and reduced expression of genes encoding receptor subunits, ion channels, and other signal transduction components (**Figure 3H-J**). Importantly, many of these direct HSF1 targets also exhibit altered HSF1 occupancy in HD mice (**Figure 2E**), suggesting that dysregulation of HSF1-dependent transcriptional programs may contribute to disease-associated synaptic dysfunction.

Finally, when we examined HSF1 direct target genes that are predominantly involved in formation and stabilization of the postsynaptic density and actin remodeling, which are uniquely occupied in WT mice but not in HD mice (**Figure 2E,H**), we found that several of these genes showed significant differential expression in *Hsf1*^+/-^ mice (highlighted in bold in the heatmap), while many additional targets exhibited consistent trends toward altered transcript levels (**Figure 3K**). Among the top five HSF1 direct synaptic targets upregulated in *Hsf1*^+/-^ mice we found *Cadm1* (Cell adhesion molecule 1/ SynCAM1), *Sorcs3* (Sortilin-related VPS10 domain containing receptor 3) and *Csmd2* (CUB and Sushi multiple domain 2) all of which participate in synaptic organization/adhesion and stabilization^83–85^. On the contrary, *Oprd1* (opioid receptor delta 1) was the topmost downregulated gene among the HSF1 direct synaptic targets. Altogether, these findings demonstrate that loss of HSF1 induces increased expression of structural synaptic genes associated with synapse organization and maintenance, while it simultaneously reduces expression of key neuromodulatory receptors. Moreover, the broad transcriptional changes observed following HSF1 reduction extend beyond direct HSF1 targets, indicating the presence of downstream regulatory networks through which HSF1 may indirectly influence synaptic gene expression. These downstream effectors likely represent additional mechanisms by which HSF1 contributes to the maintenance of synaptic programs (**Supplementary Figure 5**)

### Chronic depletion of systemic HSF1 exacerbates age-related transcriptional dysregulation

The transcriptional alterations observed following HSF1 reduction in young mice raised the possibility that HSF1 may also function as a critical regulator of synaptic gene programs during aging. We conducted and compared RNA-seq data between 2-month-old (young) and 12-month-old WT and *Hsf1*^+/-^ (middle-aged) striata (**Figure 4A, B, Supplementary File 2**). Comparison of young and middle-aged WT striatum identified a relatively restricted transcriptional signature (139 DEGs: 28 downregulated and 113 upregulated), characterized predominantly by enrichment of immune and glial activation pathways (**Figure 4C, Supplementary Figure 6A-C**). Dysregulation in 12-month-old WT aligned with canonical markers of striatal aging^86^, primarily an upregulation in transcripts related to phagocytosis (*Clec7a*), complement pathways (*C3*), and apoptosis (*Pmaip1*). Interestingly, middle-aged mice (12 months) also showed enhanced expression of genes associated with synaptic function like the cell adhesion molecule binding *Neo1* and actin binding *Tprn* compared to young mice (2-months) (**Figure 4D, Supplementary Figure 6A-C**). No significant GO categories were found among the downregulated genes. When comparing middle-aged and young *Hsf1*^+/-^, we found an exacerbated transcriptional dysregulation (**Figure 4B, E**) affecting 2,934 genes (1,471 genes down-regulated and 1,463 genes up-regulated) (**Figure 4B, E**). Importantly, GO cellular components demonstrated that the most significantly impacted cellular components in *Hsf1*^+/-^ middle-aged mice were enriched in synaptic transcripts (**Supplementary Figure 6D-I**). In addition, GO analysis showed that genes related to synaptic function and axon guidance were split into both up-regulated and down-regulated subsets, while genes related to oxidative stress were significantly upregulated and genes related to extracellular matrix were significantly downregulated (**Figure 4D**). Dysregulation of genes in these categories was exacerbated in middle-aged *Hsf1*^+/-^ mice compared to WT (**Figure 4F**). This data indicates that deficits in HSF1 exacerbate normal age-related transcriptional phenotypes and potentially also accelerate the weakening of synaptic connections associated with aging by modulating levels of transcripts important for synaptic function and axonal growth.

**Figure 4.**
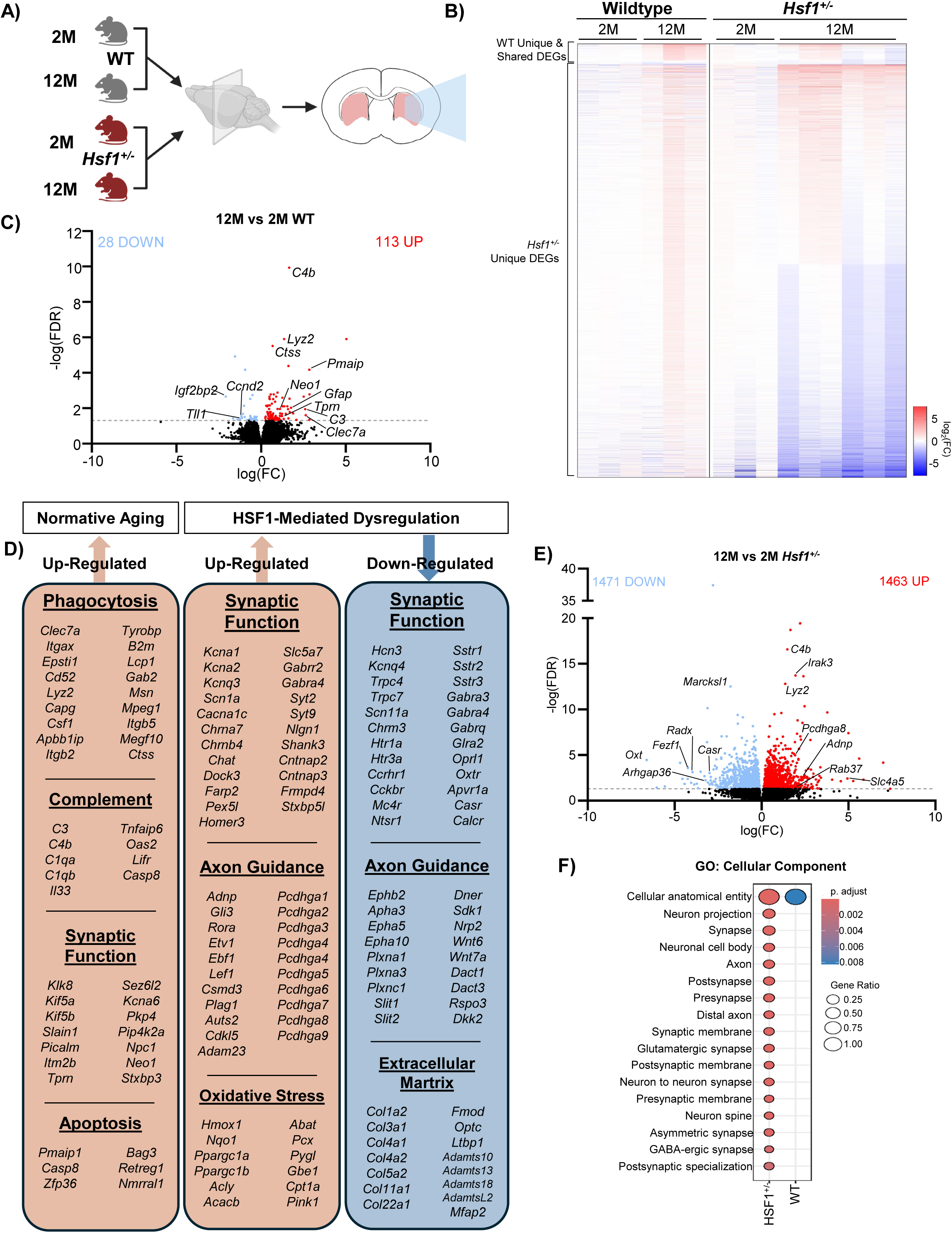
Chronic depletion of systemic HSF1 exacerbates age-related transcriptional dysregulation. **(A)** Schematic of RNA-seq experiment in 2-month- and 12-month-old WT and *Hsf1^+/-^* mice. **(B)** Heat map of DEGs in WT and *Hsf1^+/-^* mice at 2 and 12 months of age. Transcript levels are relativized to 2-month-old group for both wildtype and *Hsf1^+/-^* mice. **(C)** Volcano plot of DEGs from WT 12-month-old vs 2 month-old. **(D)** Gene lists for enriched biological functions in normative aging (12-month-old vs. 2-month) in WT and *Hsf1^+/-^* mice. DEGs: q<0.05. n=3-4 mice/group. **(E)** Volcano plot of DEGs from *Hsf1^+/-^* 12-month-old vs. 2-month-old. **(F)** Gene ontology for cellular component for DEGs between 2-month-old and 12-month-old striatum from WT and *Hsf1^+/-^* mice. DEGs: q<0.05 determined by DESeq2 (v1.46.0). n=3-6 mice/group.

### Neuronal HSF1 deletion remodels the proteomic composition of synaptic, cell adhesion, and actin-binding proteins altering shaft synapses

Our findings that HSF1 deficiency selectively disrupts T-S synapses across multiple conditions (**Figure 1**), together with the identification of HSF1 binding to and regulation of synaptic genes (**Figures 2 and 3**), demonstrate that HSF1 is a key regulator of synaptic homeostasis in the mammalian CNS. We therefore next investigated how HSF1 deficiency alters the synaptoproteome of striatal neurons. To better assess the influence of HSF1 over the synaptoproteome, we restricted the deletion of HSF1 to striatal neurons (HSF1-cKO) (**Figure 5A, Supplementary Figure 1H-K**). We isolated striatal synaptosomes and performed quality control experiments via immunoblotting (**Figure 5B**) and transmission electron microscopy (**Figure 5C**). Initial quantitative liquid chromatography–mass spectrometry analysis revealed a total of 3,223 identified proteins with a balanced distribution of synaptic proteins: 25.7% localized predominantly to presynaptic compartments, 32.0% to postsynaptic compartments, and 29.9% detected at both pre- and postsynaptic sites (**Figure 5D**). Importantly, nearly a third of the total identified proteins (968 proteins) were also verified using the SynGO bioinformatics tool^87^ (**Figure 5E**).

**Figure 5.**
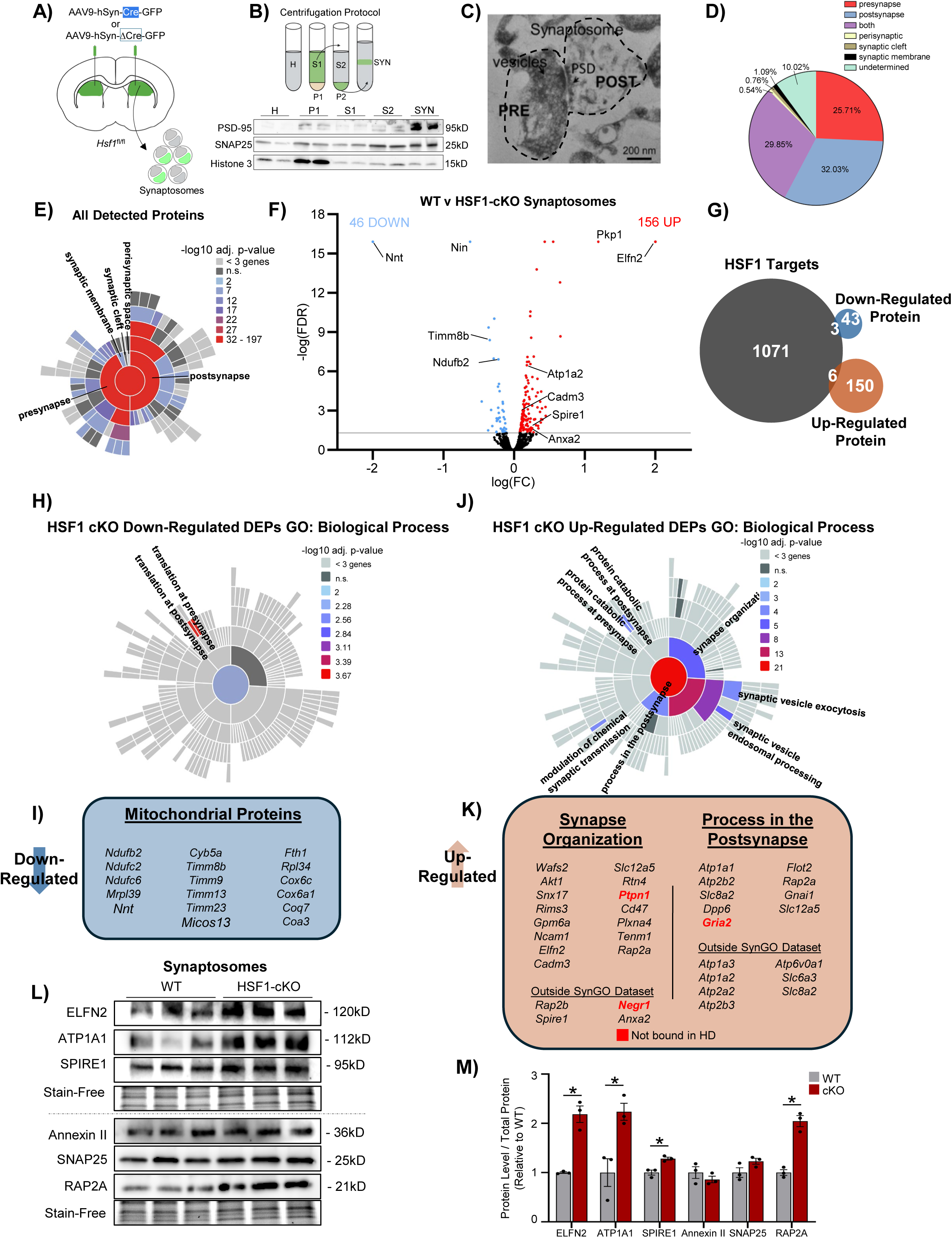
Synaptosomes from HSF1-cKO present increased accumulation of proteins involved in post-synaptic organization and decreased mitochondrial proteins. **(A)** Schematic of intracerebral injection of AAV9 viruses expressing Cre-recombinase or ΔCre-recombinase into adult 6-month-old striatum in HSF1^fl/fl^ mice (HSF1-cKO) mice. **(B)** Top: Schematic of synaptosome isolation pipeline. Bottom: Immunoblotting of synaptic proteins PSD95/SNAP25 and nuclear protein Histone H3 from synaptosome fractions. Homogenate (H), P1 (Pellet fraction 1), S1 (soluble fraction 1), S2 (soluble fraction 2), SYN (synaptosome fraction. **(C)** Transmission electron micrograph of synaptosome preparations. Scale Bar 200nm. **(D)** Relative percentages of cellular component represented across all proteins detected in synaptosome fractions by mass spectrometry. **(E)** SynGO analysis sunburst for cellular component for all proteins detected in synaptosome fractions. **(F)** Volcano plot of differentially expressed proteins (DEPs) in synaptosomes isolated from WT and HSF1-cKO striatum. **(G)** Venn diagram of DEPs in 6-month-old HSF1-cKO striatum and genes detected in 2-month-old WT HSF1 ChIP-seq. **(H)** SynGO analysis sunburst for biological process for proteins that are down-regulated in HSF1-cKO striatum. **(I)** Representative list of mitochondrial genes encoding for down-regulated DEPs. **(J)** SynGO analysis sunburst for biological process for proteins that are down-regulated in HSF1-cKO striatum. **(K)** Representative lists of genes detected in the gene ontology categories synapse organization and process in the postsynapse encoding up-regulated DEPs. **(L)** Immunoblotting of up-regulated DEPs detected in HSF1-cKO mice. **(M)** Quantification of western blot L using ImageJ. DEPs: p<0.05 determined with Proteome Discoverer. n = 4 synaptosome preparations/group. Two-Tailed Student’s t-test (M) * p<0.05. Error bars ±SEM.

We then compared the striatal synaptoproteome between WT and HSF1-cKO and found a total of 202 differentially expressed proteins (DEPs). 46 proteins were significantly down-regulated and 156 proteins were significantly up-regulated (**Figure 5F**). We cross-referenced the list of genes occupied by HSF1 and the DEPs obtained from synaptosome analyses (**Figure 5G**). We identified three direct HSF1 targets that were significantly up-regulated: LRPAP1, GNG7, NPLOC4, and six direct HSF1 targets that were significantly down-regulated: TMEM43, CEP350, IFIT3, PTPN1, GluR2, NEGR1. Across all DEPs, down-regulated DEPs were strongly enriched in mitochondrial proteins and proteins involved in local protein synthesis (**Figure 5H,I; Supplementary Figure 7A-D**). However, further analyses assessing total neuronal mitochondrial density in HSF1-cKO mice or oxygen consumption rate (OCR) measurements in cell or mouse models lacking HSF1 did not detect significant changes between genotypes or conditions **(Supplementary Figure 7E-M**).

Up-regulated proteins in HSF1-cKO synaptosomes were primarily enriched in processes across the pre- and postsynaptic compartment with two major categories: synapse organization and process in the postsynapse (**Figure 5J, K, Supplementary Figure 8**) of which *Ptpn1* and *Gria2* were direct HSF1 targets and are differentially bound in HD mice. A subset of upregulated proteins was validated via immunoblotting from isolated synaptosomes (**Figure 5L, M**). Interestingly, we noticed that several proteins within the upregulated synapse organization category were also associated with actin binding functions such as the formation and stabilization of actin patches (i.e. WAFS2, SPIRE1, ANXA2, RAP2A, RAP2B, GPM6A, CADM3, and NEGR1) of which *Negr1* is a direct HSF1 target that is differentially bound in HD **(Figure 2E; Supplementary File 1)**. Actin patches are accumulations of actin fibers within the dendritic shaft that are integral to the formation of the postsynaptic density at shaft synapses^88^. Previous reports have identified that T-S synapses make ∼25% of their total connections onto the dendritic shaft of striatal neurons, while <1% of C-S synapses make shaft connections^73^. We therefore hypothesized that HSF1-dependent alterations in the synaptoproteome, particularly in proteins involved in cell adhesion and actin dynamics, selectively influence the structural organization of shaft synapses, thereby explaining the specific role of HSF1 in regulating T-S synapse density.

We assessed whether HSF1 deficiency specifically alters shaft synapses and whether this is related to their architectural dysregulation. We utilized super-resolution microscopy with the Nikon NSPARC system in HSF1-cKD mice (**Figure 6A-C, Supplementary Figure 9**). This model was chosen because it enables sparse neuronal labeling with cytoplasmic GFP allowing complete visualization of dendritic morphology and synaptic structures at high resolution. We then imaged C-S and T-S synapses in GFP+ dendrites by co-staining with PSD-95 and VGLUT1 or VGLUT2 respectively and classified them as occurring either on dendritic spines (axospinous synapses) or within the dendritic shaft (shaft synapses)^88^. As expected, no significant depletion of C-S axospinous synapses was detected (**Figure 6D-F**). We found a small fraction of axodendritic synapses when co-staining with GFP-VGUT1-PSD95 (<2 shaft synapses/20 μm dendrite segment) with many dendritic segments showing complete absence of shaft synapses (**Figure 6G**). However, no significant differences were found between WT and HSF1-cKD. When assessing T-S synapses at the dendritic spine or the dendritic shaft we found that the previously observed depletion of T-S synapse density upon HSF1 deficiency (**Figure 1**) was related to the selective decline in T-S shaft synapses (**Figure 6H-K**).

**Figure 6.**
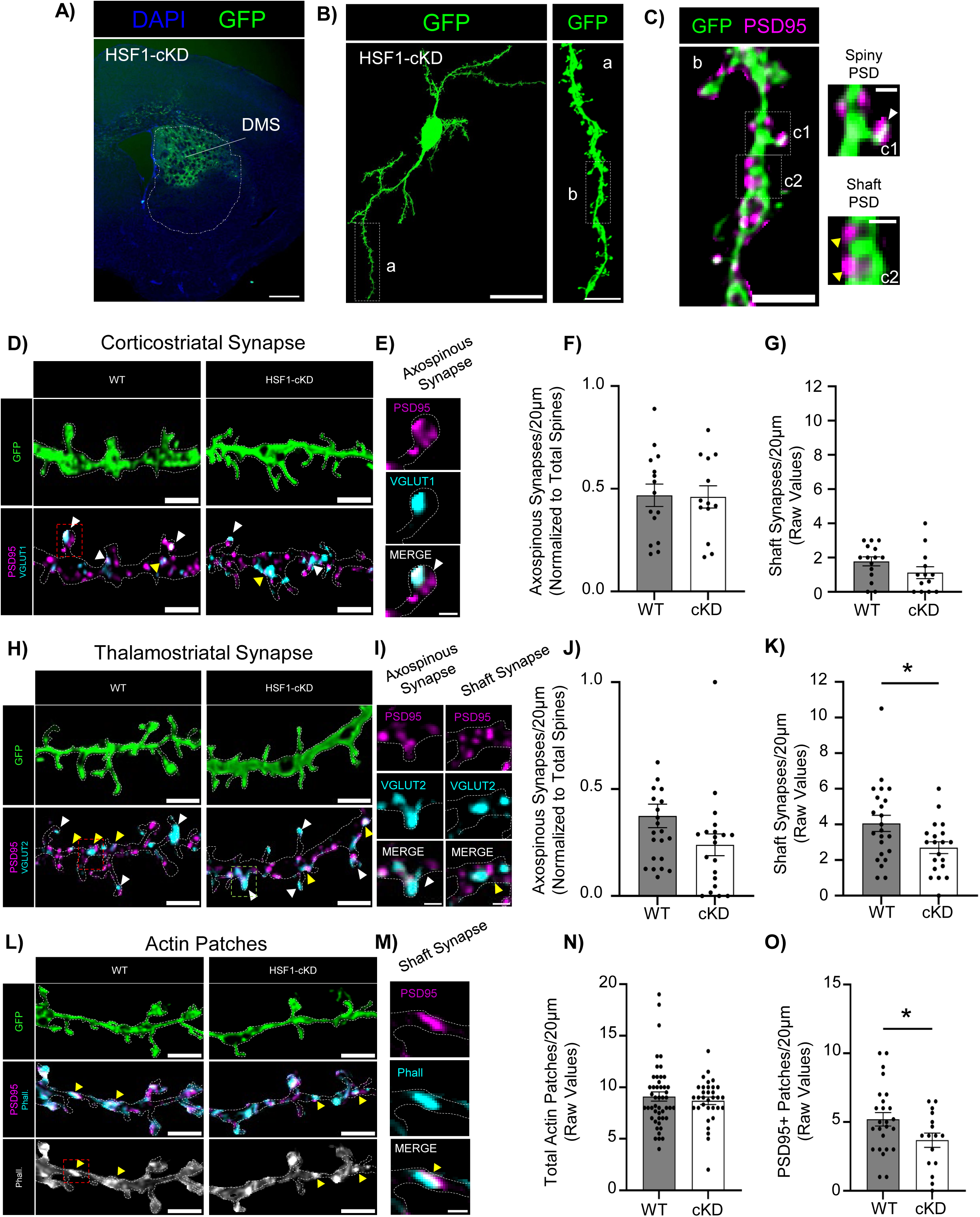
Loss of striatal HSF1 drives disrupts PSD formation within actin patches at thalamo-striatal shaft synapses. **(A)** Representative coronal section of intracranial injection of AAV9-U6-sh*Hsf1*-CMV-GFP in 6 month-old WT mice (HSF1-cKD) (Scale Bar 100 μm). **(B)** Left: Post-processed medium spiny neuron from HSF1-cKD mice (Scale Bar: 25 μm). Right: inlay GFP+ dendrite (Scale Bar: 5 μm). **(C)** Dendritic GFP+ section from B immunostained with PSD95 illustrates a representative spiny or shaft postsynaptic density (PSD) (Scale Bars: 2 μm, 500nm inlay). **(D)** Representative images of dendrites in 6-month-old striatum stained to detect corticostriatal synapses (PSD95+VGLUT1+) (Scale Bars: 2 μm). **(E)** Representative axospinous C-S synapse from D Scale Bars: 500 nm). **(F-G)** Quantification of axospinous (F) and shaft (G) corticostriatal synapses in HSF1-cKD striatum relative to total spine density in 20 μm dendritic sections. **(H)** Representative images of dendrites in 6-month-old striatum stained to detect thalamostriatal (PSD95+VGLUT2+) synapses (Scale Bars: 2 μm). **(I)** Representative axospinous (green box in H) and shaft (red box in H) T-S synapses from H (Scale Bars: 500 nm). **(J-K)** Quantification of axospinous (J) and shaft (K) thalamostriatal synapses in HSF1-cKD striatum relative to total spine density. **(L)** Representative images of dendrites in 6-month-old striatum immunostained to detect shaft synapses with actin patches (Phalloidin) (Scale Bars: 2 μm). **(M)** Representative axodendritic shaft synapse (red box in L) (Scale Bars: 500 nm). **(N-O)** Quantification of total actin patches (Phall+) (N) and shaft synapses (PSD95+/Phall) in WT and HSF1-cKD striatum. Arrows indicate axospinous synapses (white) and shaft synapses (yellow). Two-Tailed Student’s t-test (D-E,G-H,J-K). # p<0.1, * p<0.05, ** p<0.005, *** p<0.0005. n=3 mice/13-15 cells/group. 1-2 dendrites were analyzed per cell. Each point represents a 20 μm dendrite length. Error bars ±SEM.

To further connect synaptic deficits with dysfunction in actin patch regulation, we co-stained slices with the actin dye phalloidin and the postsynaptic marker PSD95 to detect high density accumulations of actin at shaft synapses^88^. We defined actin patches as high intensity phalloidin signal >1 μm in length. We did not detect any deficit in the total number of actin patches in GFP+ dendrites (**Figure 6L-N**) indicating that the initial nucleation signals that drive the formation of an actin patch are not apparently impacted by HSF1. However, when we further examined the colocalization of PSD95 and actin patches, indicative of a postsynaptic density and shaft synapse formation, we found that significantly fewer actin patches contained PSD95 (**Figure 6L,M, O**). Additionally, when we measured colocalization of actin patches with an opposing VGLUT2+ presynaptic terminal, we detected a trending decrease in colocalization (**Supplementary Figure 9C-D**). This data indicates that while actin patches may still form upon HSF1 depletion, the abnormal expression of actin binding proteins that occurs in the absence of HSF1 might contribute to impairing either the recruitment or retention of the postsynaptic machinery and alignment to presynaptic release sites. Taken together, this data strongly suggests that the specific loss of T-S synapses following loss of HSF1 is driven, at least in part, by destabilization of actin patches necessary for shaft synapses.

### HSF1 loss and disruption of T-S shaft synapses converge on deficits in cognitive flexibility

T-S projections are strongly involved in the regulation of state signaling and cognitive flexibility (CF)^32,33^. CF decline has also been reported in HD and aged mice^89–91^ where we found selective decline in T-S synapses. We therefore explored whether loss of striatal HSF1 was sufficient to drive impairments in CF. We assessed CF performance in control and HSF1-cKD mice in a two-choice visual discrimination task. In this task, mice must learn to differentiate between two images and associate interaction with a touchscreen displaying one image with the administration of a reward. Mice were food-restricted and trained to a goal criterion of 80% correct trials across two consecutive days before moving on to a reversal phase (**Figure 7A**). In the initial discrimination phase of this task, the orbitofrontal and visual cortex along with the dorsomedial striatum are highly involved^92–94^. HSF1-cKD mice showed no significant difference in the total number of days to reach criteria (**Figure 7B, C**). Additionally, HSF1-cKD mice did not display impairment in their learning rate, defined as the slope of linear regression for performance across trial days (**Figure 7D**). This indicated that discrimination learning was not affected in HSF1-cKD mice. However, when mice transitioned into a reversal phase in which the previously unrewarded image is now rewarded, and for which the dorsomedial and thalamic regions are more involved^95,96^, HSF1-cKD mice needed significantly more days than their control counterparts to reach criteria (**Figure 7E, F**). Once again, HSF1-cKD mice showed no deficit in overall learning rate during the reversal phase, (**Figure 7G**) indicating that this deficit is driven primarily by perseverative behaviors in early trial days consistent with impairment in overcoming consolidated, habitual motor behaviors.

**Figure 7.**
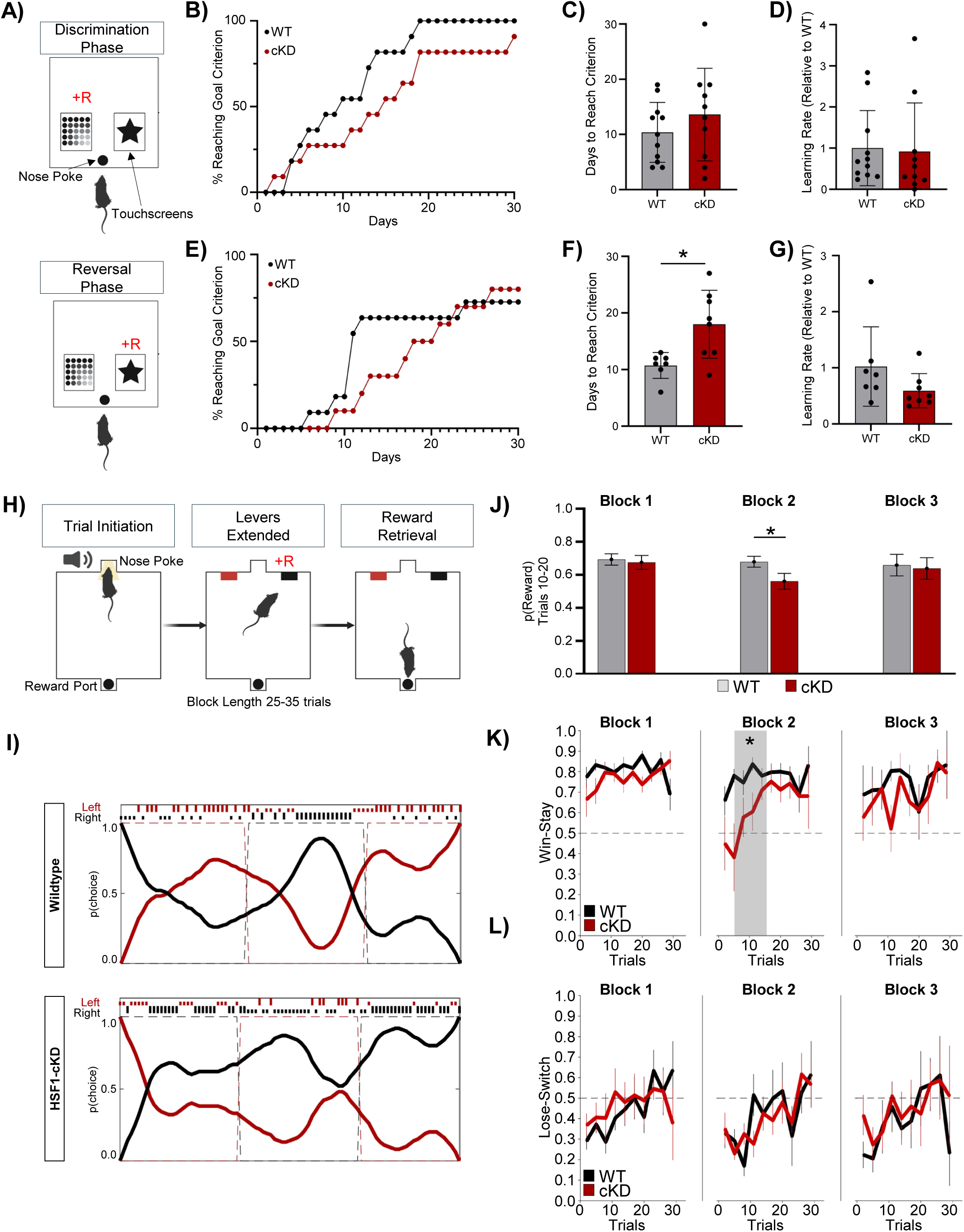
Depletion of striatal HSF1 impairs cognitive flexibility. **(A)** Schematic of two-choice visual discrimination task (2-VD) used in HSF1-cKD mice (n=10-11 mice/group). **(B)** Tracking data of -percentage of mice to reach goal criterion (80% correct trials, 2 consecutive days) during discrimination phase. **(C)** Days to reach goal criterion in discrimination phase. **(D)** Learning rate of mice during discrimination phase calculated as slope of linear regression for daily performance across all trial days. **(E)** Tracking data of percentage of mice to reach goal criterion (80% correct trials, 2 consecutive days) during reversal phase. **(F)** Days to reach goal criterion in reversal phase. **(G)** Learning rate of mice during reversal phase calculated as slope of linear regression for daily performance across all trial days. **(H)** Schematic of deterministic lever press task used in HSF1-cKD mice (n=6 mice/group). **(I)** Representative performance graphs from one hour long session in control scramble (WT) and HSF1-cKD mouse. Full tick marks indicate rewarded lever choice. Half tick marks indicate unrewarded lever choice. Dotted lines track reward contingency for each lever (0% or 100% rewarded). Plotted curves show relative probability of choice for each lever. **(J)** Probability of reward across 3 blocks during deterministic lever press task. **(K)** Win-stay behavior across 3 blocks during deterministic lever press task. **(L)** Lose-switch behavior across 3 blocks during deterministic lever press task. Two-Tailed Student’s t-test (C,D,F,G,J). Wilcoxon Rank Test (K,L) * p<0.05. Error bars ±SEM.

We next asked whether this impairment in reversal learning was unique to highly consolidated behaviors. To do this, we utilized a deterministic lever press task in which mice entered randomized block lengths of 25-35 trials across a 1-hour long session in which one lever within the operant chamber is always rewarded and the other is not. After completion of a block, the reward contingencies for each lever are switched. In this way, mice experience multiple reversal events over the course of a single training day and avoid consolidation of reward contingency to a single lever (**Figure 7H**). Similar to what we observed in the two-choice visual discrimination task, HSF1-cKD mice obtained fewer rewards than sh-SCR controls specifically during the second Block, when the reward contingency reversed to the previously unrewarded lever (p(reward) at trials 10–20: 0.68 ± 0.03 vs 0.56 ± 0.04), with no difference in Blocks 1 or 3 **(Figure 7J)**. Because the behavioral task is deterministic, the rewards achieved on each trial across blocks are a direct readout of choice quality. To determine specific behavioral deficits leading to a decreased likelihood of attaining rewards, we examined the pattern of two choice strategies that indicate how animals change their behavior to reward outcomes. Win-stay behavior, the probability of repeating a choice on the next trial if it yielded a reward, was significantly reduced in HSF1-cKD mice also selectively during Block 2 (0.77 ± 0.03 vs 0.61 ± 0.05) **(Figure 7K)**, whereas lose-switch behavior, the probability of abandoning a choice that is unrewarded, was unaffected across the two groups in every block (Block 2: 0.23 ± 0.02 vs 0.26 ± 0.02) **(Figure 7L)**. This behavioral strategy dissociation indicates that HSF1-cKD mice detect and abandon unrewarded actions normally, but fail to consolidate a newly rewarded action once it is discovered. These data point to an impaired reward credit assignment selective to unexpected positive outcomes as a driver of a broader perseverative deficit. This illustrates that these mice show perseverative behavior immediately following switch points during this task that cannot be attributed to habitual behaviors and are instead driven by deficits in cognitive flexibility. During both the two-choice visual discrimination task and deterministic lever press task, HSF1-cKD mice demonstrated similar task engagement in total trials completed per session (**Supplementary Figure 10A-D**). In addition, we detected no difference in latency to lever press or entry to reward port (**Supplementary Figure 10E,F**), which we have used as a proxy for motivation to perform the task.

To determine whether conditional depletion of striatal HSF1 selectively alters CF or if instead affects a wide range of behaviors we assessed motor performance and spatial memory in HSF1-cKD. Intriguingly, HSF1-cKD mice performed significantly more correct alternations in the spontaneous Y-maze, which could be interpreted as further evidence for a bias towards repetitive movements similar to the perseveration observed after switch points (**Supplementary Figure 10G,H)**. HSF1-cKD mice traveled similar distances as their control counterparts in the open field (**Supplementary Figure 10I,J**) and demonstrated equivalent motor learning in the accelerating rotarod test across trial days (**Supplementary Figure 10K,L**). Lastly, we performed a battery of additional behavioral tasks in HSF1-cKO mice and confirmed similar effects as in HSF1-cKD (**Supplementary Figure 10M-Q**). Taken together, these data demonstrate that the loss of striatal HSF1 is sufficient to cause selective behavioral impairments affecting CF, presumably caused by the HSF1-mediated disruption of synaptic organization and T-S shaft synapses.

## Discussion

The goal of this study was to understand how HSF1 regulates T-S synapses and influences cognitive functions that depend on T-S synapse stabilization. Through our analysis, we have identified a novel role of HSF1 in the mammalian central nervous system, under both disease and physiological conditions, that consists of the regulation of key structural synaptic genes and proteins that are necessary for the stabilization of T-S synapses that has implications in the regulation of cognitive flexibility.

Previous studies have established HSF1 as a versatile transcription factor that regulates distinct transcriptional programs in response to cancer, metabolic perturbations, and cellular stress^42,76,97–100^. However, HSF1’s regulatory functions in the central nervous system have remained poorly defined. Here, we provide the first comprehensive characterization of HSF1-dependent regulatory programs in the mammalian brain by integrating physiological and disease contexts with genome-wide DNA binding, transcriptional profiling, proteomic and molecular analyses, and behavioral assessments. This integrated approach uncovered previously unrecognized roles of HSF1 in the regulation of brain function under both normal and pathological conditions. Our observation that HSF1 interacts with over 1000 gene targets during critical points of striatum maturation highlights the broad implications of HSF1 in the regulation of postnatal neurodevelopment, synapse formation, and neuronal signaling. This data is supported by observations in primary hippocampal neuronal cultures from *Hsf1*^-/-^ mice showing impaired spinogenesis^65^. Our data is also supported by early HSF1 ChIP-seq studies in immortalized striatal ST*Hdh* control (Q7) and HD (Q111) cells revealing that HSF1 targets upon stress are not directly associated with proteostasis, but with cytoskeletal binding, focal adhesion and GTPase activity^82^, which are gene categories identified in our Chip-seq analyses. Importantly, the integration of our ChIP-seq, RNA-seq, and synaptoproteomic analyses has also uncovered a previously unrecognized role for HSF1 in regulating postsynaptic specialization and synaptic stability. This function was not identified in earlier studies using cultured ST*Hdh* cells which lacked the complex synaptic architecture and activity-dependent neuronal interactions necessary to reveal context-dependent regulation of synaptic genes and proteins. In line with our data, studies in primary hippocampal neurons treated with an Hsp90 inhibitor (17-AAG), which results in the activation of HSF1, connected HSF1 activity with the regulation of BDNF, PSD-95, synapsin I, and synaptophysin^50^. Our prior studies also showed HSF1 can directly bind to and regulate members of the PSD gene family in both mice and human brain tissues^62^. Collectively, these findings consolidate the key role of HSF1 in the regulation of synaptic architecture and synaptic stability.

Decline in the levels and/or activity of HSF1 have been widely reported in several neurological conditions including HD and normative aging^48,49,101–103^. Here we showed that HSF1 genome-wide occupancy, especially on genes involved in synaptic organization and specialization, is dramatically altered in early HD, even when the levels of HSF1 have not robustly declined yet^49^. This suggests an early dysfunction of HSF1 in HD that precedes HSF1 degradation and impairs its ability to bind to concrete regions of the chromatin. HSF1 DNA binding is regulated by several processes including posttranslational modifications and protein-protein interactions^42,82,104–106^. One such protein-protein interaction is with the chromatin remodeler CTCF^82^. The enrichment of a CTCF-like motif at HSF1 occupied loci raises the possibility that HSF1 cooperates with chromatin architectural proteins to regulate neuronal gene expression. Because CTCF organizes chromatin loops and topologically associated domains, HSF1 recruitment to CTCF-associated loci could facilitate the coordinated regulation of genes involved in synaptic function by integrating stimulus-dependent transcription with higher-order genome organization. One caveat to this hypothesis is the lack of changes in the magnitude of HSF1-CTCF interaction reported between control and HD conditions^82^, implying that other regulatory proteins that are dysregulated in HD could also be responsible for the impairment in HSF1 DNA-binding. Our data further suggests that the inability of HSF1 to bind to and regulate these HSF1-mediated synaptic signatures contributes to the T-S synapse decline observed in HD.

Although HSF1 is classically viewed as a transcriptional activator of heat shock proteins^99,100,107^, it is now well established that HSF1 can also function as a direct or indirect transcriptional repressor in a highly context-dependent manner^108–110^. Our data demonstrated that lack of HSF1 resulted in the significant upregulation of several genes and proteins, especially those associated with synaptic functions and cytoskeletal/actin binding, while other unrelated genes were downregulated. This implies HSF1 acts as a transcriptional repressor of synaptic and actin binding genes. However, not all genes occupied by HSF1 were differentially expressed following HSF1 depletion, and conversely, many DEGs were not identified as direct HSF1 targets. This mismatch between HSF1 chromatin occupancy and the broader transcriptional response suggests that loss of HSF1 produces widespread secondary effects on neuronal gene expression that is extrapolated to the synaptic proteome. These indirect effects may be mediated, at least in part, through HSF1-dependent regulation of other transcriptional regulators and signaling intermediates, including p53, HIF-1α, and MYC^111–113^.

An important question is how the observed regulatory effects of HSF1 influence T-S synapses. Despite some recent controversy^6^, it is widely accepted that HD mice exhibit an early and selective loss of T-S synapses^38,72,114,115^ and we previously demonstrated that T-S synapse density is closely linked to HSF1 levels and/or activity^49,62^. Specifically, genetic stabilization of HSF1 in HD mice through deletion of the kinase CK2α’, which promotes HSF1 degradation in HD, rescued T-S synapse density^49^. Conversely, chronic depletion of HSF1 in control mice was sufficient to induce selective T-S synapse loss^62^, demonstrating that HSF1 is both necessary and sufficient for the maintenance of these synapses. We have also confirmed that conditional depletion of HSF1 in the striatum with a non-cell selective or a neuronal-specific approach is also sufficient to reduce T-S synapse density. These results suggested that prior effects seen in *Hsf1*^+/-^ or *Hsf1*^-/-^ mice were not exclusive to neurodevelopmental problems and that HSF1 exerts a direct role over synapse regulation also in the adult brain. In addition, we extended these findings by showing that T-S synapses also undergo a selective age-dependent decline in the healthy striatum, paralleling the progressive reduction in HSF1 levels^62^. Our obtained data indicates that HSF1 regulates essential postsynaptic specialization and actin binding proteins whose coordinated and balanced expression is necessary for the appropriate formation of shaft T-S synapses. A surprising result was the observation that many of these proteins were upregulated upon HSF1 depletion. We believe this result calls into question the simplistic understanding that down-regulation of synaptic proteins drives synaptic loss and up-regulation indicates formation or rescue. The dynamic nature of synapses, which undergo near constant restructuring in the form of spine growth, insertion and removal of postsynaptic receptors, protein and vesicular trafficking, exo- and endocytosis of vesicles, and many others, requires a finely tuned concert of transcriptional/translational control to mediate these processes. Therefore, dysregulation, whether in loss or gain of protein concentrations, may tip the balance of these dynamics resulting in synaptic dysfunction and disassembly. Importantly, we found increased levels of proteins involved in actin patch assembly and maintenance, more specifically in actin nucleation signaling proteins (WAVE2, Spire-1, Rap2A, Rap2B), actin branching, anchoring of receptors, and alignment to the presynaptic terminal (Annexin II, CADM3, NEGR1, M6A). The dysregulation of these proteins did not impair the initial formation of actin patches, however, the presence of postsynaptic machinery inside of the actin patch was significantly depleted. On the contrary, because axospinous synapses do not rely on the complex formation of actin patches, they were not affected. This finding suggests that either the recruitment or maintenance of essential postsynaptic proteins is impaired within the actin patch and specifically affects T-S synapses. We hypothesize that this is driven by changes in cytoskeletal dynamics that selectively weaken the postsynaptic density within shaft synapses.

Interestingly, we also found a significant depletion in mitochondrial proteins from HSF1-cKO synaptosomes. There is substantial evidence that HSF1 regulates mitochondrial function and biogenesis in different contexts, including HD^49,58,116,117^. Indeed, HSF1 directly binds to and regulates the peroxisome proliferator co-activator PGC-1α (*Ppargc1a*), the master transcriptional coactivator of mitochondrial biogenesis and oxidative metabolism^118,119^. However, our studies assessing total mitochondrial density via immunofluorescence or mitochondrial function in cells and mice lacking HSF1 measuring OCR did not show substantial mitochondrial deficits upon HSF1 depletion. An important nuance here is that our studies only reflect whole cell mitochondrial parameters and not local dendritic mitochondria. Distinctions in respiratory function between mitochondria resident to the synapse and mitochondria found in other cellular compartments has been previously reported^120–122^. Because our measurements do not account for this distinction, we cannot rule out that synaptic mitochondria may be uniquely affected in HSF1 cKO mice. Additionally, depletions in mitochondrial proteins from HSF1-cKO synaptosomes could be driven by dysfunctions in mitochondrial trafficking or retention within the synaptic compartment, processes that depend on actin-based anchoring mechanisms^123–125^. Accordingly, dysregulation of HSF1-dependent actin-binding proteins may compromise the efficient retention of mitochondria within shaft synapses, resulting in reduced synaptic mitochondrial content in parallel with the diminished retention of postsynaptic elements that we have reported.

A major function of the T-S circuit is the regulation of CF^23,32,33,35,36^. Consistent with this role, pharmacological inactivation of the parafascicular thalamus (Pf), which projects to the dorsomedial striatum (DMS), impairs reversal learning without affecting initial acquisition^36^. Likewise, disruption of the Pf–DMS pathway through anatomical lesions or pathway-specific inhibition preserves initial goal-directed learning but results in persistent deficits in updating action–outcome associations following changes in behavioral contingencies^33–35^. Given the marked depletion of T-S synapses following HSF1 deletion, we investigated whether HSF1 contributes to CF. We found that the loss of striatal HSF1 is sufficient to drive selective impairment in CF without affecting initial acquisition or other behaviors like motor performance and spatial memory. Mice with systemic deletion of *Hsf1* exhibit a range of behavioral abnormalities that become more pronounced with age, including reduced exploratory behavior, impaired working memory, and progressive motor deficits^126,127^. To our knowledge, this is the first study to identify cognitive flexibility as a behavioral function regulated by HSF1, suggesting that distinct HSF1-dependent neural circuits underline different behavioral domains. However, our study does not establish a direct causal link between HSF1-mediated T-S synapse loss, dysregulation of actin-binding and synaptic protein expression, and impaired CF. Therefore, future experiments combining circuit-specific manipulation of T-S projections with behavioral testing in HSF1-cKO mice will be required to directly test this relationship.

## Acknowledgments

We thank Dr. Ezquiel Marron Fernandez Velasco and the University of Minnesota Viral Vector and Cloning Core for their assistance in producing the viruses used in this study. We thank Dr. Erin Lind and the University of Minnesota Mouse Behavior Core for their guidance and expertise in conducting behavioral studies. We also thank Dr. Chengkai Dai for his generous gift of the *Hsf1*^fl/fl^ mouse model used in this study and Dr. Yun You for his assistance in *in vitro* fertilization to produce the *Hsf1^+/-^* mouse line. We also thank Dr. Brian Freeman for his generous gift of reagents and assistance with the EMSA experiments in this study. This work was supported in part by the resources and staff at the University of Minnesota University Imaging Centers (UIC; SCR_020997) and the University of Minnesota Center for Metabolomics and Proteomics (CMSP). The Orbitrap Eclipse instrumentation platform used in this work was purchased through High-end Instrumentation Grant S10OD028717 from the NIH.

## Author Contributions

Conceptualization—RG-P; Methodology—NBR, NZ, RM, PG, KG, AB; Formal analysis—NBR, KG, YZ, AH, AB; Investigation, RG-P, NBR, NZ, RM, PG, KG; Writing—original draft presentation, NBR; writing—review and editing, RG-P, NBR; Funding acquisition—RG-P. All authors have read and agreed to the published version of the manuscript.

## Institutional Review Board Statement

All animal work was conducted according to procedures approved by the University of Minnesota Institutional Animal Care and Use Committee (IACUC) in compliance with the National Institute of Health guidelines for the care and use of laboratory animals under the approved animal protocol 2605-43924A.

## Data Availability Statement

All data generated in this study is presented in the current manuscript. Datasets have been deposited into the following public repositories: ChIP-seq—GEO (accession# GSE342107), RNA-seq—GEO (accession# GSE342108). All other data is available upon request from the corresponding author.

## Supplementary Figure Captions

**Supplementary Figure 1. Confirmation of thalamostriatal synapse loss with Homer1 as postsynaptic marker and in HSF1-cKO in striatal neurons. (A)** Western blot of HSF1 levels in 3-month-old wildtype and HD and 18-month-old wildtype striatal protein extracts (n=3 mice/group). **(B)** Quantification of HSF1 protein levels from A. **(C)** Representative images of C-S (VGLUT1+) and T-S (VGLUT2+) synapses in the striatum of 3-month-old HD and wildtype mice (n=3 mice/3 slices/group). **(D)** Quantification of C-S synapses from C. **(E)** Quantification of T-S synapses from E. **(F)** Western blot from ST*Hdh*^Q^^7/Q^^7^ cells infected with increasing concentrations of sh*Hsf1* or shScr (control). **(G)** Representative HSF1 signal in striatal regions infected with AAV9-U6-sh*Hsf1*-CMV-GFP and regions that are not virus-infected. **(H)** Western blot of HSF1 protein levels in HSF1-cKO striatum (AAV9-hSyn-Cre-GFP) and control (AAV9-hSyn-ΔCre-GFP) (n=6 mice/group). **(I)** Quantification of HSF1 protein levels from G. (J) *Hsf1* mRNA transcript levels from HSF1-cKO and control striatum (n=6 mice/group). **(K)** Representative coronal section of viral spread of AAV9-hSyn-Cre-GFP intracranial injection and HSF1 protein levels in infected striatal neurons. **(L)** Representative images of C-S (VGLUT1+) and T-S (VGLUT2+) synapses in the striatum of 6-month-old HSF1-cKO and control (n=3 mice/3 slices/group). **(M)** Quantification of C-S synapses from K. **(N)** Quantification of T-S synapses from K. Scale Bars: 5 μm. One-Way ANOVA (A) Two-Tailed Student’s t-test (D,E,H,I,L,M). * p<0.05, **p<0.005. Error bars ±SEM.

**Supplementary Figure 2. Gene ontology of ChIP-seq datasets. (A)** Gene ontology enrichment for cellular component of genes uniquely bound by HSF1 in wildtype striatum at 1-month. **(B)** Gene ontology enrichment for biological process of genes uniquely bound by HSF1 in wildtype striatum at 1-month. **(C)** Gene ontology enrichment for molecular function of genes uniquely bound by HSF1 in wildtype striatum at 1-month. **(D)** Gene ontology enrichment for cellular component of genes uniquely bound by HSF1 in wildtype striatum at 2-months. **(E)** Gene ontology enrichment for biological process of genes uniquely bound by HSF1 in wildtype striatum at 2-months. **(F)** Gene ontology enrichment for biological process of genes uniquely bound by HSF1 in HD striatum at 2-months. **(G)** Gene ontology enrichment for biological process of genes bound by HSF1 across 1-month-old wildtype and 2-month-old wildtype and HD striatum. **(H)** Gene ontology enrichment for molecular function of genes bound by HSF1 across 1-month-old wildtype and 2-month-old wildtype and HD striatum. **(I)** Gene ontology enrichment for biological process of genes bound by HSF1 across 1-month-old and 2-month-old wildtype striatum, but not HD. **(J)** Gene ontology enrichment for molecular function of genes bound by HSF1 across 1-month-old and 2-month-old wildtype striatum, but not HD. Only significantly enriched gene ontology categories shown. Categories not included for a given gene list indicate no significant enrichment detected. Gene enrichment analysis performed using ShinyGo bioinformatics tool.

**Supplementary Figure 3. Validation of differential HSF1 binding to target genes in HD and abundance of binding motifs. (A)** Chromatin immunoprecipitation for *Gria2* binding site detected in ChIP-seq in ST*Hdh*^Q^^7/Q^^7^ and ST*Hdh*^Q^^111/Q^^111^ cells. **(B)** Chromatin immunoprecipitation for *Oprd1* binding site detected in ChIP-seq in ST*Hdh*^Q^^7/Q^^7^ and ST*Hdh*^Q^^111/Q^^111^ cells. (n=3-5 cultures/group) **(C)** Total peaks detected in 1-month-old wildtype and 2-month-old wildtype and HD striatum for top DNA-binding motifs identified through *de novo* motif enrichment in ChIP-seq. **(D)** Percentage of total peaks that coincide with top DNA-binding motifs identified in ChIP-seq in 1-month-old wildtype and 2-month-old wildtype and HD striatum. Student t-test (A,B). # p<0.1, * p<0.05.

**Supplementary Figure 4. Gene ontology of HSF1-mediated 2-month-old transcriptional dysregulation. (A)** Gene ontology enrichment for cellular component of up-regulated genes in 2-month-old *Hsf1^+/-^* striatum. **(B)** Gene ontology enrichment for biological process of genes up-regulated genes in 2-month-old *Hsf1^+/-^* striatum. **(C)** Gene ontology enrichment for molecular function of genes up-regulated genes in 2-month-old *Hsf1^+/-^* striatum. **(D)** Gene ontology enrichment for cellular component of genes down-regulated genes in 2-month-old *Hsf1^+/-^*striatum. **(E)** Gene ontology enrichment for biological process of genes down-regulated genes in 2-month-old *Hsf1^+/-^* striatum. **(F)** Gene ontology enrichment for molecular function of genes down-regulated genes in 2-month-old *Hsf1^+/-^* striatum. Only significantly enriched gene ontology categories shown. Categories not included for a given gene list indicate no significant enrichment detected. Gene enrichment analysis performed using ShinyGo bioinformatics tool.

**Supplementary Figure 5. Ingenuity pathway analysis (IPA) predicts downstream effectors of transcriptional and translational dysregulation after loss of HSF1. (A)** IPA for top 30 upstream regulators of DEGs from 2-month-old wildtype or *Hsf1^+/-^* RNA-seq dataset. **(B)** Proposed signaling cascade of effector proteins driving transcriptional and protein dysregulation. **(C)** Lists of DEPs from synaptoproteome of 6-month-old control or HSF1-cKO mice alongside proposed transcription regulators identified using ingenuity pathway analysis. **(D)** IPA for top 30 upstream regulators of DEPs from 6-month-old HSF1-cKO synaptosomes.

**Supplementary Figure 6. Gene ontology of age-related transcriptional dysregulation in wildtype and *Hsf1^+/-^* mice. (A)** Gene ontology enrichment for cellular component of up-regulated genes in 12-month-old wildtype striatum. **(B)** Gene ontology enrichment for biological process of genes up-regulated genes in 12-month-old wildtype striatum. **(C)** Gene ontology enrichment for molecular function of genes up-regulated genes in 12-month-old wildtype striatum. **(D)** Gene ontology enrichment for cellular component of genes up-regulated genes in 12-month-old *Hsf1^+/-^* striatum. **(E)** Gene ontology enrichment for biological process of genes up-regulated genes in 12-month-old *Hsf1^+/-^* striatum. **(F)** Gene ontology enrichment for molecular function of genes up-regulated genes in 12-month-old *Hsf1^+/-^* striatum. **(G)** Gene ontology enrichment for cellular component of genes down-regulated genes in 12-month-old *Hsf1^+/-^*striatum. **(H)** Gene ontology enrichment for biological process of genes down-regulated genes in 12-month-old *Hsf1^+/-^* striatum. **(F)** Gene ontology enrichment for molecular function of genes down-regulated genes in 12-month-old *Hsf1^+/-^* striatum. Only significantly enriched gene ontology categories shown. Categories not included for a given gene list indicate no significant enrichment detected. Gene enrichment analysis performed using ShinyGo bioinformatics tool.

**Supplementary Figure 7. Loss of striatal HSF1 does not affect total mitochondrial number or mitochondrial function in striatal neurons. (A)** SynGO sunburst gene ontology for cellular component of down-regulated DEPs in HSF1-cKO synaptosomes. **(B)** Gene ontology enrichment for cellular component of down-regulated DEPs in HSF1-cKO synaptosomes. **(C)** Gene ontology enrichment for biological process of down-regulated DEPs in HSF1-cKO synaptosomes. **(D)** Gene ontology enrichment for molecular of down-regulated DEPs in HSF1-cKO synaptosomes. **(E)** Representative images of VDAC1 protein levels in post-processing medium spiny neurons in the striatum of HSF1-cKO and control mice. **(F)** Quantification of VDAC1 signal in medium spiny neurons from F. (n=3 mice/group). Each data point represents a single cell. **(G)** Oxygen consumption rate in ST*Hdh*^Q^^7/Q^^7^ striatal cell line between wildtype or ST*Hdh*:*Hsf1*^+/-^ cells. (n=11 cultures/group) **(H)** Oxygen consumption rate from bulk striatal lysates of 12-month-old wildtype or *Hsf1^+/-^* mice. **(I)** Reactive oxygen species production from bulk striatal lysates of 12-month-old wildtype or *Hsf1^+/-^* mice. **(J)** Complex I OXPHOS capacity. **(K)** Complex I coupling efficiency. **(L)** Complex I & II OXPHOS capacity. **(M)** Complex I & II coupling efficiency. n=7-8 mice/group. Scale Bars: 5 μm. Two-Tailed Student’s t-test. # p<0.1.

**Supplementary Figure 8. Gene ontology of up-regulated proteins from HSF1-cKO striatal synaptosomes. (A)** SynGO sunburst gene ontology for cellular component of up-regulated DEPs in HSF1-cKO synaptosomes. **(B)** Gene ontology enrichment for cellular component of up-regulated DEPs in HSF1-cKO synaptosomes. **(C)** Gene ontology enrichment for biological process of up-regulated DEPs in HSF1-cKO synaptosomes. **(D)** Gene ontology enrichment for molecular of up-regulated DEPs in HSF1-cKO synaptosomes.

**Supplementary Figure 9. HSF1-cKD in the striatum results in dendritic spine loss and disruption of T-S shaft synapses. (A)** Representative images of GFP+ dendrites in 6-month-old control and HSF1-cKD mice. **(B)** Quantification of total dendritic spines from A. **(C)** Representative images of GFP+ dendrites stained to detect T-S (VGLUT2+) shaft synapses with inlay of representative T-S shaft synapse. **(D)** Quantification of T-S shaft synapses measured by co-localization of VGLUT2 and phalloidin+ actin patch. Scale Bars: 2 μm (A,C); 500 nm (inlays). Two-Tailed Student’s t-test. * p<0.05. n=3 mice/26-30 cells (A) or 13-15 cells (B)/group. 1-2 dendrites were analyzed per cell. Each point represents a 20 μm dendrite length. Error bars ±SEM.

**Supplementary Figure 10. HSF1 deficiency in the adult brain does not impair motivation, motor performance, or spatial working memory. (A)** Mice weight during two choice visual discrimination task represented as percent original body weight. **(B)** Average trials during a session of the two choice visual discrimination task (n=10-11 mice/group (A-B)). **(C)** Mice weight during deterministic lever press task. **(D)** Average trials during a session of the deterministic lever press task. **(E)** Average time to lever choice during the deterministic lever press task. **(F)** Average time to enter reward port on rewarded trials during the deterministic lever press task (n=6 mice/group (C-F)). **(G)** Schematic of spontaneous Y-maze used in HSF1-cKD and HSF1-cKO mice. **(H)** Percent correct alternations in a Y-maze between HSF1-cKD and control mice. **(I)** Schematic of open field test used in HSF1-cKD and HSF1-cKO mice. **(J)** Distance traveled in 5-minute increments in an open field test between HSF1-cKD and control mice. **(K)** Schematic of accelerating rotarod test used in HSF1-cKD mice. **(L)** Accelerating rotarod test performance measured by latency to fall across 5 trial days (n=6 mice/group). **(M)** Percent correct alternations in a Y-maze between HSF1-cKO and control mice. **(N)** Distance traveled in 5-minute increments in an open field test between HSF1-cKO and control mice. **(O)** Schematic of Barnes circular maze. **(P)** Time to find escape hole across trial days between HSF1-cKO and control mice. **(Q)** Time spent in target quadrant during probe phase. 12-15 mice/group (G-O) Two-Tailed Student’s t-test. (B,D,E,F,H,Q) RM-ANOVA (A,C,J,L,N,P) *p<0.05. Error bars ±SEM.

**Supplementary Table 1:** Antibodies Used in ChIP, Immunoblotting, and Immunofluorescence.

| **Primary Antibodies** |  |  |  |  |
| --- | --- | --- | --- | --- |
| **Target** | **Use** | **Manufacturer** | **Catalog#** | **Dilution** |
| Annexin II | WB | Santa Cruz | sc-28385 | 1:1000 |
| ATP1A1 | WB | Invitrogen | PA5-110490 | 1:1000 |
| DARPP32 | IF | R&D Systems | MAB4230 | 1:500 |
| ELFN2 | WB | Novus | NBP1-90569 | 1:1000 |
| GFP | IF | Abcam | AB13970-1001 | 1:1000 |
| Histone 3 | WB | Cell Signaling Technology | 9715 | 1:1000 |
| Homer1 | IF | Synaptic Systems | 160003 | 1:200 |
| HSF1 | WB/ChIP | Bethyl | A303-176A | 1:1000 |
| HSF1 | IF | Enzo | ADI-SPA-950-F | 1:500 |
| Mouse IgG | ChIP | Cell Signaling | 5415S |  |
| PSD95 | IF | ThermoFisher | 51-6900 | 1:500 |
| PSD95 | WB | Novus | NB300-556 | 1:1000 |
| RAP2A | WB | Proteintech | 13789-1-AP | 1:1000 |
| SNAP25 | WB | Proteintech | 14903-1-AP | 1:10,000 |
| SPIRE1 | WB | Proteintech | 11295-1-AP | 1:500 |
| VDAC1 | IF | Abcam | AB15895 | 1:500 |
| VGLUT1 | IF | MilliporeSigma | AB5905 | 1:500 |
| VGLUT2 | IF | MilliporeSigma | AB2251-I | 1:1000 |
| **Secondary Antibodies** |  |  |  |  |
| **Target** | **Use** | **Manufacturer** | **Catalog#** | **Dilution** |
| AlexaFluor Goat $\alpha$ Chicken 488 | IF | Invitrogen | A-11039 | 1:200 |
| AlexaFluor Goat $\alpha$ Guinea Pig 488 | IF | Invitrogen | A-11073 | 1:200 |
| AlexaFluor Goat $\alpha$ Rabbit 594 | IF | Invitrogen | A-11012 | 1:200 |
| AlexaFluor Goat $\alpha$ Guinea Pig 647 | IF | Invitrogen | A-21450 | 1:200 |
| AlexaFluor Goat $\alpha$ Rat 647 | IF | Invitrogen | A-21247 | 1:200 |
| AlexaFluor Phalloidin 647 | IF | Invitrogen | A-30107 | 1:400 |
| Donkey $\alpha$ Rabbit IgG HRP | WB | Cytiva | NA934V | 1:5000 |
| Sheep $\alpha$ Mouse IgG HRP | WB | Cytiva | NXA931V | 1:5000 |

## Materials and Methods

### Mouse Stains

For this study, all mouse strains were bred on the C57BL/6J or C57BL/6N background. We used the heterozygous zQ175 knock-in mouse model obtained from Jackson Labs (Stock No. 027410; Bar Harbor, ME, USA). Sperm from HSF1 heterozygous knock-out mice (B6N(Cg)-Hsf1^tm1^(KOMP)^Vlcg^ / JMmucd) was obtained from the Mutant Mouse Resource and Research Center (University of California, Davis; Davis, CA, USA; Stock No. 048101-UCD) and generated by the Knockout Mouse Phenotyping Program (KOMP^2^). *In vitro* fertilization using C57BL/6N females was conducted by the Mouse Genetics Laboratory at University of Minnesota. *Hsf1^+/-^*^:tm1^ mice were crossbred with CMV-Cre mice (B6.C-Tg(CMV-cre)1Cgn/J (Stock No. 006054) to delete the Neomycin cassette flanked by loxP sites (*Hsf1*^+/-:tim1.1:Cre^). The *Hsf1*^+/-:tim1.1:Cre^ line was crossed with C57BL/6N to remove the Cre gene (*Hsf1*^+/-:tm1.1^, referenced as *Hsf1^+/-^* in this study). *Hsf1*^fl/fl^ mice were obtained as a generous gift from Dr. Chengkai Dai at the National Institutes of Health National Cancer Institute^128^. All animal care and sacrifice procedures were approved by the University of Minnesota Institutional Animal Care and Use Committee (IACUC) in compliance with the National Institutes of Health guideline for the care and use of laboratory animals under the approved animal protocol 2605-43924A.

### AAV intracranial injections

shRNA against HSF1 was obtained from Sigma MISSION TRC1 lentiviral human shRNA library. Non-targeting sequence shScr plasmid (pLKO.1-TRC Cloning Vector) or shHSF1 plasmid (TRCN0000007480, Sigma) were previously validated in murine cells^82^. U6-shScr and U6-shHSF1 were subcloned into AAV9-CMV-GFP (AAV9-U6-sh*Hsf1*-CMV-GFP and AAV9-U6-shScr-CMV-GFP) and were packaged by the University of Minnesota Viral Vector Core. AAV9-hSyn-Cre-GFP and AAV9-hSyn-ΔCre-GFP were provided by the Stanford University Gene Vector and Virus Core. During surgical procedure, mice were anesthetized using gaseous isoflurane (up to 5%) and maintained on a heating pad at 37°C. Injections were performed using a Stoelting digital mouse stereotaxic instrument (Stoelting Co., Wood Dale, IL, USA). Burr holes were drilled into the skull at injection sites and administered using a Hamilton syringe (33G, 1” needle, point type 2) at +0.5 A/P, ±2.3 M/L, -3.1 D/V relative to bregma. Total injection volume was 0.5 µl administered at 0.1 µl/min. Wounds were sutured closed and removed 10 days post-injection. Carprofen (20mg/kg) was administered for analgesia prior to surgery and for 3 days post-surgery. All analyses were conducted at least 4 weeks post-injection.

### Cell Lines

Mammalian cell lines used in this study were the mouse-derived striatal cells ST*Hdh*^Q7/Q7^ and ST*Hdh*^Q111/Q111^ (Coriell Cell Repositories; Camden, NJ, USA). ST*Hdh*:*Hsf1*^+/-^ cells were generated using pSpCas9-(BB)-2A-GFP (PX458, Addgene #48138; Watertown, MA, USA) and a gRNA targeting *Hsf1*-exon9 (5′-caccGAGTACCCGAGGGCTGTGAGGCT CATGGGCTCCCGACACTCCcaaa-3′). Generation of ST*Hdh*:*Hsf1^+/-^* cells was described previously.^62^ Cells were grown at 33°C in Dulbecco’s modified Eagle’s medium (DMEM, Genesee; El Cajon, CA, USA) supplemented with 10% fetal bovine serum (FBS), 100 U mL^−1^ penicillin/streptomycin, and 100 μg mL^−1^ G418 (Gibco, Thermo Fisher Scientific; Waltham, MA, USA).

### Mouse Sample Preparations

Mice were pericardially perfused with tris-buffered saline (TBS) (25 mM Tris-Base, 135 mM NaCl, 3 mM KCl, pH7.6) supplemented with 7.5 μM heparin. Brains were removed from the skull and one hemisphere was microdissected to remove the striatum which was used for downstream immuoblotting or RT-qPCR analysis. The other hemisphere was drop fixed in 4% paraformaldehyde (PFA) in TBS for 4 days at 4°C before transferring the brains to 30% sucrose in TBS until fully impregnated and then embedded in a 2:1 mixture of 30% sucrose in TBS:OCT (Tissue-Tek) and stored at -80°C. Brains were cryo-sectioned at 16 μm and stored in 50:50 Glycerol:TBS at -20°C for long term storage.

### Immunofluorescence, Microscopy, and Synapse Density Analysis

For standard immunofluorescent experiments, brain sections were blocked in 5% normal goat serum (NGS) in TBST (0.3% Triton) and probed with primary antibodies in 5% NGS in TBST overnight at 4°C. The following day sections were incubated with secondary antibodies in 5% NGS in TBST for 1 hour. Slides were mounted in ProLong Gold with DAPI (Invitrogen). Images were acquired on a Leica Stellaris 9 confocal microscope running Leica LAS-X image software with a 10X (NA 0.4), 20X (NA 0.75), or 63X (NA 1.4) lens. Confocal scans were acquired at a pixel resolution of 512 x 512. Maximum projections were generated and data was analyzed using FIJI ImageJ image analysis software.

For slices used in synapse density analysis, three independent coronal brain sections were used for each mouse containing dorsal striatum (bregma 0.5-1.1 mm). Slices were blocked in 20% NGS in TBST and probed with primary antibodies in 10% NGS in TBST overnight at 4°C overnight. The following day sections were incubated with secondary antibodies in 10% TBST for 2h. Slides were mounted in ProLong Fold with DAPI (Invitrogen). Images were acquired on a Leica Stellaris 9 confocal microscope running Leica LAS-X image software with a 63X lens (NA. Confocal scans (optical section depth 0.34 mm, 15 sections per scan) were acquired at a pixel resolution of 856 x 856. Maximum projections of 3 optical scans per slice were generated. Synapse density analysis was performed blinded using the PunctaAnalyzer Plugin (Durham, NC, USA) on ImageJ as previously described^49^. A minimum of three slices per animal and three animals per genotype and time point were analyzed.

For slices used in super-resolution studies, sections were stained as above for synapse density analyses. Images were acquired on a Nikon Eclipse Ti2 with NSPARC detector running Nikon Elements image software with a 100X lens (NA 1.45). Confocal scans optical section depth (0.15 μm) were acquired at a pixel resolution of 2048 x 2048. Maximum projections were generated and analyzed using Imaris software. Briefly, GFP signal was used to define surfaces and signal outside of GFP+ surfaces was masked. Imaris spot detection was used to define VGLUT1/2+ (0.4 μm; PSF 0.8 μm), PSD95+ (0.25 μm; PSF 0.5 μm), and phalloidin (1 μm; PSF 2 μm) puncta. Co-localization of defined puncta was determined using a MATLAB distance transformation Imaris plugin from PSD95+ puncta. Synapses were defined as co-localization of <250nm for VGLUT1/2:PSD95 and VGLUT2:phalloidin and 0 nm for PSD95:phalloidin. Categorization of spiny and shaft synapses was performed manually and blinded to group. Information on all antibodies used can be found in **Supplementary Table 1.**

### Immunoblot Analysis

Sample preparation and immunoblotting conditions were performed as previously described^49^. Cell protein extracts were prepared in cell lysis buffer (25 mM Tris pH 7.4, 150 mM NaCl, 1 mM EDTA, 1% Triton X-100, 0.1% SDS) supplemented with HALT phosphatase and protease inhibitors (ThermoScientific; Waltham, MA, USA). Striatal protein extracts from one hemisphere of mice were prepared in tissue lysis buffer (2 5mM Tris pH 7.4, 150 mM NaCl, 1 mM EDTA, 1% Triton X-100, 1% SDS) supplemented with HALT phosphatase and protease inhibitors (ThermoScientific; Waltham, MA, USA). Protein samples were separated on 4-20% TGX Stain-Free gels (BioRad; Hercules, CA, USA) and transferred to a nitrocellulose membrane (BioRad 0.2 μm. Membranes were blocked in 5% dry milk in TBST (Tween) for 1 hour before probing with primary antibodies (**Supplementary Table 1**) in 2.5% dry milk in TBST overnight at 4°C. Membranes were probed with secondary HRP-tagged antibodies **(Supplementary Table 1)** in 2.5% dry milk in TBST for 1 hour and then imaged using SuperSignal Western Blot Substrate Kit Pico PLUS or Femto PLUS using an Amersham ImageQuant 800 (Cytiva; Marlborough, MA, USA). Quantitative analyses were performed using ImageJ software and normalized to GAPDH controls.

### Chromatin Immunoprecipitation Sequencing

15 mg of frozen striatal tissue was cross-linked with 37% formaldehyde for 15 minutes at 4°C followed by glycine quenching (125 mM final concentration) for 5min. Tissue was pelleted (5,000rpm for 5 minutes) before homogenizing in ice-cold PBS. Tissue was centrifuged at 2,000 x *g* for 7 minutes ay 4°C, resuspended in 1X Buffer A (Cell Signaling; Danvers, MA, USA) supplemented with DTT and HALT protease and phosphatase inhibitors and incubated at 4°C for 10 minutes. Tissue was centrifuged again (2,000 x *g*; 5 minutes; 4°C) and resuspended in 1X Buffer B (Cell Signaling; Danvers, MA, USA) supplemented with micrococcal nuclease and DTT. Lysates were centrifuged (16,000 x *g*; 1 minute; 4°C) and resuspended in immunoprecipitation buffer (IPB) (50 mM Tris pH 7.5, 150 mM NaCl, 1 mM EDTA, 1% Triton X-100) and sonicated three times for 20s at 40% amplitude. Samples were centrifuged at 9,400 x *g* for 10 minutes at 4°C. An aliquot (30µl) was saved for control input DNA. 2µg of anti-HSF1 (Bethyl) or anti-IgG was added and samples were incubated overnight at 4°C. Dynabeads^TM^ Protein G (Invitrogen; Waltham, MA, USA) were added and incubated 4 hours at 4°C. Immunocomplexes were washed, eluted using elution buffer (10 mM Tris-HCl, pH 8, 1 mM EDTA, pH 8, 1% SDS) and crosslinking was reversed at 65°C for 12 hours. Protein was digested by addition of Proteinase K and incubated at 37°C for 1.5 hours. Chromatin was purified using the Qiaquick min-elute PCR purification kit (Qiagen; Germantown, MD, USA) per the manufacturer’s instructions. Antibodies used in immunoprecipitation can be found in **Supplementary Table 1.** ChIP-seq was performed by University of Minnesota Genomics Center. The purified DNA samples were quantified using a fluorimetric PicoGreen assay and ran on Agilent Tapestation (Agilent; Santa Clara, CA, USA) to assess sizing. Library preparation was performed using the ThruPLEX DNA-Seq Kit (Takara Bio; San Jose, CA, USA) according to manufacturers instructions. Pooled libraries were denatured and diluted to the appropriate clustering concentration. The libraries were then loaded onto a NovaSeq paired end flow cell (Illumina; San Diego, CA, USA) and clustering occurs on board the instrument. Once clustering is complete, the sequencing reaction immediately begins using Illumina’s 2-color SBS chemistry. Upon completion of read 1, two separate 8 or 10 base pair index reads were performed. Finally, the clustered library fragments were re-synthesized in the reverse direction thus producing the template for paired end read 2. Base call (.bcl) files for each cycle of sequencing were generated by Illumina Real Time Analysis (RTA) software. The base call files and run folders were streamed to servers maintained at the Minnesota Supercomputing Institute. Primary analysis and de-multiplexing were performed using Illumina’s bcl2fastq v2.20 For analysis, raw paired-end ChIP-seq reads in fastq format were first assessed for base call quality, cycle uniformity, and contamination using FastQC. Raw reads were mapped to the mouse reference genome (ensemble GRCm39) via Burrows-Wheeler Aligner (BWA)^129^. Duplicate samples were merged into a single file, excluding one wild type 1 month sample which failed the QC check. Binding peaks were called with Model-Based Analysis for ChIP-Seq version 2.0 (MASC2)^130^ using default parameters, and deepTools^131^ was used for data visualization. Motif exploration tools, MEME^80^ was used to identify enriched motifs and PWMScan^132^ was used to scan for enrichment of known motifs. ChIP-AP (v2023)^133^ was used for functional annotation of ChIP-seq peaks. Genes were grouped given their proximity to specific peaks, such as peaks in wildtype 2 month sample but not in HD. Enriched pathways were analyzed using the ShinyGo bioinformatics toolset^77^. HSF1 binding was confirmed in ST*Hdh*^Q7/Q7^and ST*Hdh*^Q111/Q111^ using RT-qPCR with the following primers: *Gria2* Forward: 5’ - GCCCATAGTATGCACGGATCA - 3’ and Reverse: 5’ - AGAGCAAAACAGGCCTAGCA - 3’, Oprd1 Forward: 5’ - GGACACCCGTTGCTTTGTTC - 3’ and Reverse: 5’ - TGTCTCCCAAGCCCTAAGGT - 3’

### Electrophoretic Mobility Shift Assay

Recombinant human HSF1 protein (2.5 ng; Abcam ab78795) and IR800-labelled DNA probes (2.5 ng) (IDT; Coralville, IA, USA) were incubated together for 30 minutes at 37°C in binding buffer (12 mM HEPES, 12% glycerol, 0.06 M KCl, 0.3 M EDTA, 0.6 mM DTT) and then placed on ice for 30 minutes. Samples not containing HSF1 were supplemented with BSA (2.5 ng). Samples were then diluted 5X with binding buffer and 20ul of sample was loaded and separated on a 4-20% TGX Stain-Free gels (BioRad; Hercules, CA, USA) under native conditions. Gels were imaged on a LI-COR Odyssey CLX. Positive binding was measured by the presence of a band shift. The following nucleotide sequences were used to confirm HSF1 binding via EMSA: *Gria2* Forward: 5’ - IR800-CTG CCT CTG CCT CTG - 3’ and Reverse: 5’ - CAG AGG CAG AGG CAG - 3’, *Oprd1* Forward: 5’ - IR800-TGT AAT CCC AGC ATC - 3’ and Reverse: 5’ - GAT GCT GGG ATT ACA - 3’, *Fmnl3* Forward: 5’ - IR800-AGT TCC GGG ACA GCC - 3’ and Reverse: 5’ - GGC TGT CCC GGA ACT - 3’.

### RNA-Sequencing

RNA-sequencing analysis was performed by the University of Minnesota Genomics Center. RNA samples were prepared using the RNeasy Plus Universal Mini Kit (Qiagen; Germantown, MD, USA). We created 32 unique dual-indexed (UDI) TruSeq stranded mRNA libraries. The libraries were combined in a single pool and sequenced on a NovaSeq Charter Service S4. All expected barcodes were detected and mean depth for the libraries was ≥20 million reads. Mean quality scores for all libraries were ≥Q30. Libraries were gel size selected to have inserts of ∼200 bp. Differential genes expression was determined with DESeq2 (v1.46.0) using default settings^134^. Genes with a FDR ≤ 0.05 were considered significant. Outliers’ identification was performed using Cook’s distance (DESeq2). Enriched pathways were analyzed using the ShinyGo bioinformatics toolset^77^.

### RNA-Preparation and RT-qPCR

RNA was extracted from mouse striatal tissues using the RNeasy extraction kit (Qiagen; Germantown, MD, USA) according to the manufacturer’s instructions. cDNA was prepared using the Superscript III First Strand Synthesis System (Invitrogen; Waltham, MA, USA) according to the manufacturer’s instructions. SYBR green-based qPCR was performed using SYBR mix (Genesee Scientific; El Cajon, CA, USA) using a LightCycler 480 System (Roche; Basel, Switzerland). Primers used are as follows: *Gapdh* Forward: 5’ - ACACATTGGGGGTAGGAACA - 3’ and Reverse: 5’ - AACTTTGGCATTGTGGAAGG - 3’, *Hsf1* Forward: 5’ - AAGTTGTCAACAAGCTCATTCAG - 3’ and Reverse: 5’ - TCAAAGAGCACAGGCAGCTCACT - 3’, *Cdhr2* Forward: 5’ - CCCTTGTGTGTTTGCGGAAG - 3’ and Reverse: 5’ - TAGCAGCAGTTGGAGCTGTC - 3’, *Pcdhga2* Forward: 5’ - GGCGCAAAGAAATTCAACCCA - 3’ and Reverse: 5’ - TCCGTATCTCCCAGGGTCTG - 3’

### Synaptosome Isolation

Synaptosome isolation was adapted from previously published protocols^135,136^. Isolation was performed using pooled freshly dissected whole striata from two mice. Striatal tissue was gently homogenized using mechanical trituration in isomolar buffer (1 mM HEPES, 0.32M sucrose) before centrifuging in a benchtop centrifuge at 1000 x *g* for 5 minutes to pellet large cellular debris and compartments. The supernatant was then centrifuged at 12,600 x *g* for 8 minutes and the pellet was resuspended in isomolar buffer and layered on top of a isomolar buffer Ficoll gradient (Top: 7.5% Ficoll; Bottom 13% Ficoll). The gradient was centrifuged at 50,000 x *g* to separate synaptosomes. Synaptosomes were then filtered and captured on a PVDF membrane (Durapore; 0.1 μm pore), collected in tubes, and flash frozen in liquid nitrogen before being stored at -80°C until use. Synaptosome protein extracts used in western blot analysis were extracted from membranes using tissue lysis buffer on a thermal mixer (1500rpm at 60°C).

### Transmission Electron Microscopy

Synaptosomes were pelleted and fixed using 1% glutaraldehyde and 2% PFA (Electron Microscopy Sciences) in 1X PBS solution and kept at 4°C. Fixed synaptosomes were treated with 1% osmium tetroxide in PBS for 1h, washed in water then 50% and 70% ethanol before incubation in 1% uranyl acetate/70% ethanol for 1h and dehydrated in increasing alcohol concentration. Pellets were embedded in EPON and ultrathin sections were cut and mounted on a TEM grid. Samples were imaged using a JEM-1400Plus to confirm the presence of intact synaptosomes in preparations.

### Liquid Chromatography Mass Spectrometry

Proteomic analyses were performed by the University of Minnesota Center for Metabolomics and Proteomics. Synaptosome samples were acetone precipitated followed by reconstitution in 6 μl denaturing buffer (7 M urea, 2 M thiourea, 0.4 M tris-HCl ph 8, 20% acetonitrile, 10 mM TCEP, 40 mM chloroacetamide). The samples were incubated at 37°C for 30min. After incubation, the samples were centrifuged at 13000 x *g* for 10 minutes at room temperature. A Bradford assay was performed to determine protein concentration. Aliquots for each sample were made for proteolytic digestion. All the samples were brough to the same volume with denaturing buffer and then diluted 4X with water. Trypsin was added to each sample in a 1:40 enzyme to total protein ratio. Samples were incubated 16 hours at 37°C. After incubation, samples were acidified with 10% formic acid to a final concentration of 0.3% formic acid. The trypsin aliquots were cleaned up with MCX-like STAGE tips^137^. The eluted samples were then vacuum centrifuged to dryness. The dried peptides were reconstituted in 97.9:2:0.1, H2O:acetonitrile (ACN):formic acid (FA) (load solvent) and analyzed ∼100ng of each sample by capillary LC-MS with a Thermo Fisher Scientific, Inc (Waltham, MA) Dionex UltiMate 3000 RSLCnano system on-line with an Orbitrap Eclipse mass spectrometer (Thermo Scientific, Waltham MA). We injected peptides directly in load solvent and performed gradient separation on a self-packed C18 column (Dr. Maisch GmbH ReproSil-PUR 1.9 um 120 Å C18aq, 100 um ID x 35 cm length) at 55 °C with the following profile: 5% B solvent from 0 – 2 minutes, 8% B at 2.5 minutes, 21% B at 30 minutes, 35% B at 45 minutes and 90%B at 47 minutes with a flowrate of 400 nl/min from 0 – 2 minutes and 315 nl/minute from 2.5 – 47 minutes, where solvent A was 0.1% formic acid in water and solvent B was 0.1% formic acid in ACN. We employed a data independent acquisition (DIA) workflow on the Orbitrap with the following MS parameters: ESI voltage +2.1 kV, ion transfer tube 275 °C; no internal calibration; Orbitrap MS1 scan 60k resolution in profile mode from 390 – 990 m/z with 50 msec injection time; 100% (4 x 10E5) automatic gain control (AGC); we acquired tandem MS with HCD (high energy collision dissociation) activation at a fixed collision energy of 33% and orbitrap detection at 30k resolution; the precursor isolation window was set to 12 Da with a 4 Da overlap; the number of MS2 scan events was 50 per cycle and the MS2 loop count was set to 20; the MS2 injection time was 54 msec and the automatic gain control was set to 800%.

We processed peptide tandem MS using CHIMERYS 3.0.0 (https://www.msaid.de/) spectral prediction algorithm module in Proteome Discoverer 3.1.0.638 (Thermo Scientific). The *mus musculus* Universal Proteome (UP000000589; accessed August 18, 2023) protein sequence database was downloaded from UniProt.org, merged with a common lab contaminant protein database (https://github.com/HaoGroup-ProtContLib) (55402 total protein sequences), and used as the template during data processing. The CHIMERYS database search parameters were: enzyme trypsin full specificity, 1 missed cleave site, ‘store all traces’ and ‘process as one job per file.’ We specified CAM cysteine (+57.021 Da) as a static modification and oxidation of methionine (+15.995 Da) as a dynamic modification (maximum 1 dynamic modification per peptide). We applied 1% protein and peptide False Discovery Rate (FDR) and 2 peptide minimum as filters on the protein report using the Percolator algorithm^138^ in PD. Gene ontology analyses were performed using the SynGO bioinformatics tool^87^ and the ShinyGO bioinformatics tool^77^

### Two-Choice Visual Discrimination Task

Mice were singly-housed and food-restricted for the entire duration of the two-choice visual discrimination task. Mouse weight was recorded daily to maintain 80% of starting weight. When mice fell below 80% of their starting weight, supplemental food was given in cage to ensure mouse health. Mice were habituated to operant chambers for two days before training phase began. The training phase consists of three phases: (1) one day of initial touch training in which reward is administered immediately upon interaction with a touchscreen or every 30s, (2) two days of must touch training in which rewards are only presented upon interaction with a touchscreen, and (3) three days of must initiate training in which mice must nose poke into illuminated area prior to interaction with touchscreens to receive reward. After training, mice began the full two-choice visual discrimination task. In the full task, mice are placed into the operant chamber for 1 hour to achieve as many rewards as possible. The task is structured such that mice must nose poke into the illuminated reward area to initiate a trial, interact with one of two touchscreens that display distinct images, and then enter the reward port to consume reward (vanilla or strawberry Ensure). During discrimination learning, a single touchscreen is rewarded across trial days and the side and image displayed is randomized across mice. Mice are tested to a goal criterion of 80% correct trials across a session for two consecutive days. After reaching criteria, mice advance to the reversal stage of the task in which the reward contingencies of each touchscreen are switched such that the previously unrewarded screen is now rewarded. Mice are again tested to a goal criterion of 80% correct trials across two consecutive days. Both testing phases allow for a maximum of 30 days to reach criterion. If a mouse did not reach criterion in either phase, they were removed from analysis. Data was collected using a custom LabView code.

### Deterministic Lever Press Task

Mice were singly-housed and fluid restricted for the entire duration of the deterministic lever press task. Mouse weight was recorded daily to maintain 80% of starting weight. When mice fell below 80% of their starting weight, supplemental water was given in cage to ensure mouse health. The operant chambers consists of a nose poke, two extendable/retractable levers, and a reward port. The task is structured such that mice must nose poke in an illuminated chamber to begin a trial at which point both levers extend and are illuminated, the mouse then makes a lever selection and proceeds to the reward point. Trial starts are signaled by a tone in the box and illumination of nose poke chamber. Mice were habituated to operant chambers for two days before training phase began. The training phase consists of 4 phases: (1) two days of free reward accompanied by tone, (2) reward is administered only if a nose poke occurs and mice are trained to 50% engagement across trials for two consecutive days, (3) lever training in which mice must now nose poke and press a lever to receive reward and are trained to 50% completion across trials for two consecutive days, and (4) bias removal in which only one lever is illuminated and extended and mice are trained to 50% engagement across trials for two consecutive days. Mice then begin the full task as described above. Mice performed the task for a maximum of 10 weeks dependent on training phase duration. Chamber was cleaned with 70% ethanol between individual sessions. Data was collected using a custom LabView code.

### Open Field Test

Activity chambers (20”x20”x10”, made in house) were equipped with light intensity of 250 lx. Mice were placed in the center of the chamber and their behavior was recorded using ANY-maze software (Stoelting Co., Wood Dale, IL, USA) for 30 mins. Activity chamber was cleaned with 70% ethanol between mice. Males were run before and separate from females. Analyses were performed on total locomotion in the chamber.

### Rotarod Test

Mice were in accelerating mode with a ramp of 5 to 50 rpm over 5 minutes on a UgoBasile rotarod system. Mice received 4 trials a day for 4 days with a ∼15 minute intertrial interval. The trial ended when the mouse fell off the rod, or when they completed two full 360° rotations. The latency to fall was averaged across the 4 trials each day in each animal for analysis. Apparatus was cleaned with 70% ethanol between trials. Males were run before and separate from females.

### Spontaneous Y-Maze Test

Spontaneous alternation task was used to assess spatial working memory and was performed at least 20 hours after the spatial memory task. In this task, the innate response of a mouse to a new environment was evaluated. Alternation was defined as entry into all three arms, e.g., ABC, BCA, or CAB, but not CAC. The mouse was habituated in the testing room for 15-20 mins while setting up the apparatus. The animal was then placed in one arm of the maze and was allowed to move freely in all three arms for 5 minutes. Maze was cleaned with 70% ethanol between mice. Males were run before and separate from females. Latency to exit the start arm, the number and pattern of arm choices were recorded. Behavior was recorded using ANY-maze software (Stoelting Co., Wood Dale, IL, USA).

### Barnes Circular Maze

The Barnes maze is a dry-land maze test for spatial learning and memory. It consists of a circular platform with 20 holes along the perimeter (San Diego Instruments). Spatial cues were placed on all four walls of the behavioral testing room, lit to 300 lx during testing. Mice were acclimatized to the testing room for 30 min at the beginning of each training session. Training days consisted of 4 trials per day for 4 days. Training trials ended when the subject climbed into the escape box within the goal quadrant located under 1 of the holes or when the maximum trial duration of 180 s was reached. Subjects were run in small groups of five mice or less, so that no more than 20 min passed between trials for a given animal during training. On the day following the last training trial, memory was assessed in single 90 s probe trial tests, where the target escape box in the goal quadrant was replaced with a false box cover identical to the other 19 holes, and the exploration pattern of each subject was examined. Maze was cleaned with 70% ethanol between trials. Males were run before and separate from females. Behavior was recorded using ANY-maze software (Stoelting Co., Wood Dale, IL, USA).

### Mitochondrial Oxygen Consumption Rate Measurements

Oxygen consumption rate (OCR) from cultured wildtype ST*Hdh*^Q7/Q7^ and ST*Hdh*:*Hsf1*^+/-^ cells were measured using the Resipher system (Lucid Scientific, Atlanta, GA, USA). Cells were plated in 96-well plates at a staring density of 10,000 cells and cultured at 37°C in growth media overnight to allow them to adhere. OCR measurements were made using the Lucid Lab software at 37°C for up to 24 hours. Cells were then lysed and total protein concentration was measured using Pierce BCA protein assay kit (ThermoFisher). Only the last OCR measurement was used in quantifications and measurements were normalized to total protein concentration in each well as a proxy for total cell number.

*Ex vivo* OCR and hydrogen peroxide (H_2_O_2_) production from striatal tissue were simultaneously measured using the Oroboros NextGen-O2k Fluo system (Oroboros Instruments, Innsbruck, Austria). Methods for examining OCRs and H_2_O_2_ were adapted from a previously published protocol^139^, with OCRs measured using high-resolution respirometry and H_2_O_2_ measured simultaneously by fluorometry. Daily calibration of the H_2_O_2_ signal was performed by sequential injections of 0.1 μM H₂O₂ to generate a standard curve (0-0.5 μM). Measurements were conducted in assay buffer consisting of buffer Z (containing 105 mM K-MES, 30 mM KCl, 10 mM K_2_HPO_4_, 5 mM MgCl_2_.6H_2_O, 0.5 mg/ml bovine serum albumin at pH 7.1) with 1 mM EGTA, 20 mM creatine monohydrate, 10 μM Amplex UltraRed (AmR), 1 U/mL horseradish peroxidase (HRP), and 5 U/mL superoxide dismutase (SOD) at 37°C. After acquiring the H_2_O_2_ standard curve, air calibration was performed using the DatLab protocol “O_2_-calibration air”. Mice were anesthetized using isoflurane (up to 5% inhalation), decapitated, and brains rapidly removed. Extracted brains were placed in a mouse brain matrix, and a coronal section entailing the striatum was isolated. Bilateral samples of striatal tissue were collected, and wet tissue weight recorded; the average striatum wet tissue weight was 3.6 ± 0.5 mg per mouse. Collected striatal tissue was suspended in ice-cold Buffer X (7.23 mM K_2_EGTA, 2.77 mM CaK_2_EGTA, 20 mM imidazole, 0.5 mM DTT, 20 mM taurine, 5.7 mM ATP, 14.3 mM phosphocreatine disodium salt hydrate, 6.56 mM MgCl_2_.6H_2_O, 50 mM K-MES at pH 7.1) and permeabilized with saponin (50 µg/mL) for 30 minutes, followed by three 5 minutes washes in ice-cold buffer Z at 4°C. OCRs and H_2_O_2_ production were determined by sequential addition of substrates/inhibitors using Hamilton microsyringes as follows: glutamate (G, 10 mM) + malate (M, 2 mM) (G/M), pyruvate (P, 5 mM) + malate (M, 2 mM) (P/M), ADP (2.5 mM), succinate (Suc, 10 mM), cytochrome C (CyC, 4 mM), rotenone (Rot, 0.5 µM), antimycin A (AA, 5 µM) and ascorbate (A, 2 mM) + N, N, N′, N′-tetramethyl-p-phenylenediamine (T, 0.5 mM) (A/T). Glutamate, malate and pyruvate are complex I substrates, ADP is a precursor to oxidative phosphorylation (OXPHOS), succinate is a complex II substrate, cytochrome C is an electron carrier for complex III to IV, rotenone is a complex I inhibitor, antimycin A is a complex III inhibitor and ascorbate and N, N, N′, N′-tetramethyl-p-phenylenediamine are complex IV substrates. OXPHOS capacity refers to the mitochondria’s maximum ability to produce ATP. Complex I and complex I + II OXPHOS capacities were calculated by subtracting leak state respiration (C_L_, derived from glutamate and malate addition) from complex-linked active phosphorylation respiration (C_P_; determined after addition of ADP and succinate). Coupling efficiency indicates how effectively mitochondria use oxygen, i.e., how well electrons are transferred through the electron transport chain, rather than being lost as leak or uncoupled respiration. Complex I and complex I + II coupling efficiencies were calculated using the formula 1-C_P_/C_L_. OCRs and H_2_O_2_ measurements were first calculated based on milligrams of wet brain tissue using DatLab software (Oroboros Instruments, Innsbruck, Austria).

### Statistical Analysis

Sample sizes for each experiment are specified in the relevant figure legend. Data are expressed as Mean ±SEM, analyzed for statistical significance, and displayed by Prism 9 software (GraphPad, San Diego, CA, USA) unless otherwise specified here. The accepted level of significance was p<0.05, except for RNA-seq where q<0.05 was used. Deterministic lever press task was analyzed using custom MATLAB code. Statistical analysis for ChIP-seq, RNA-seq, and LC-MS datasets are described under their respective method section.

