## Supplementary Figures 1-10 for "HSF1 controls transcriptional programs that establish thalamostriatal shaft synaptic architecture and preserve cognitive flexibility"

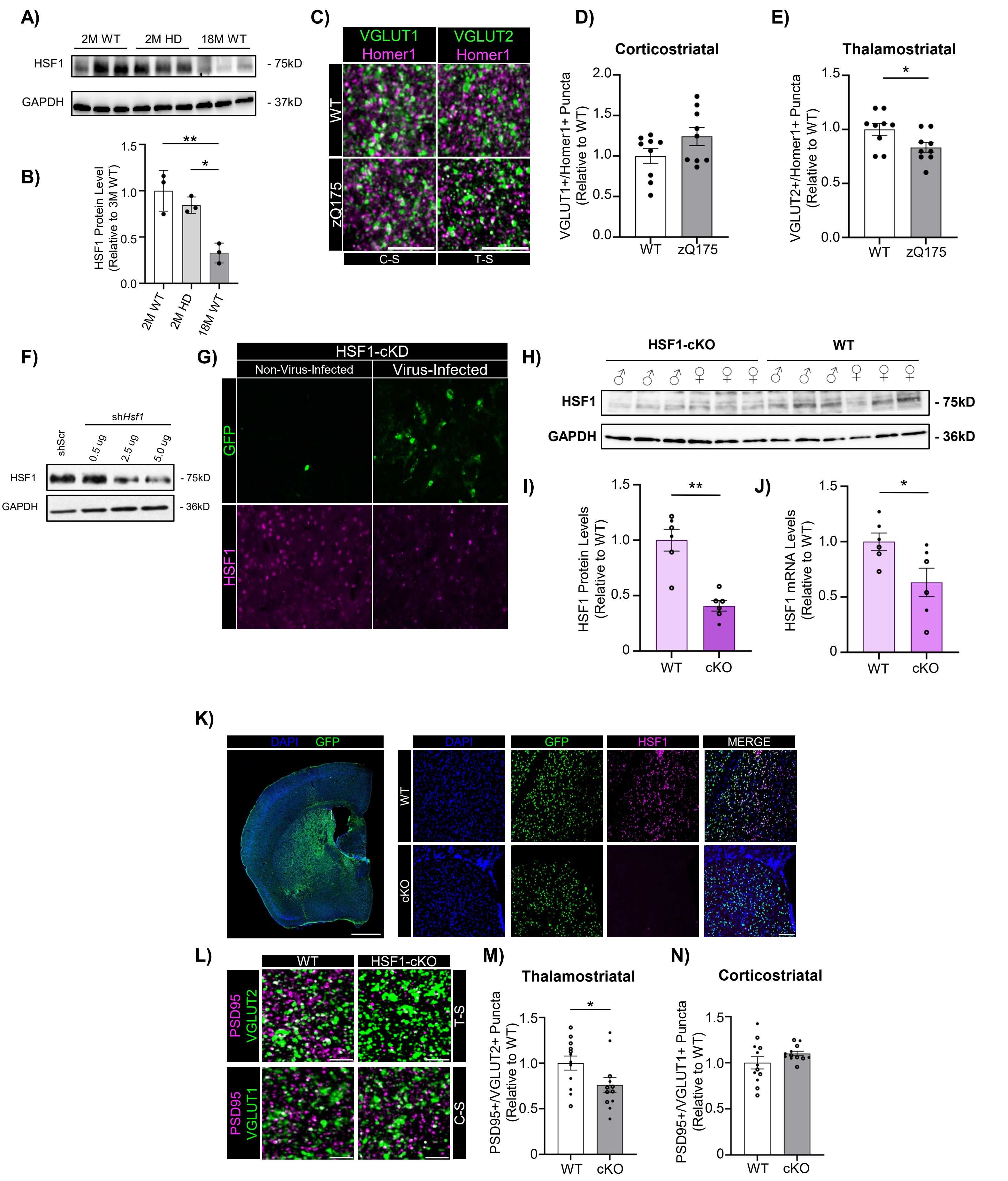


**Supplementary Figure 1**

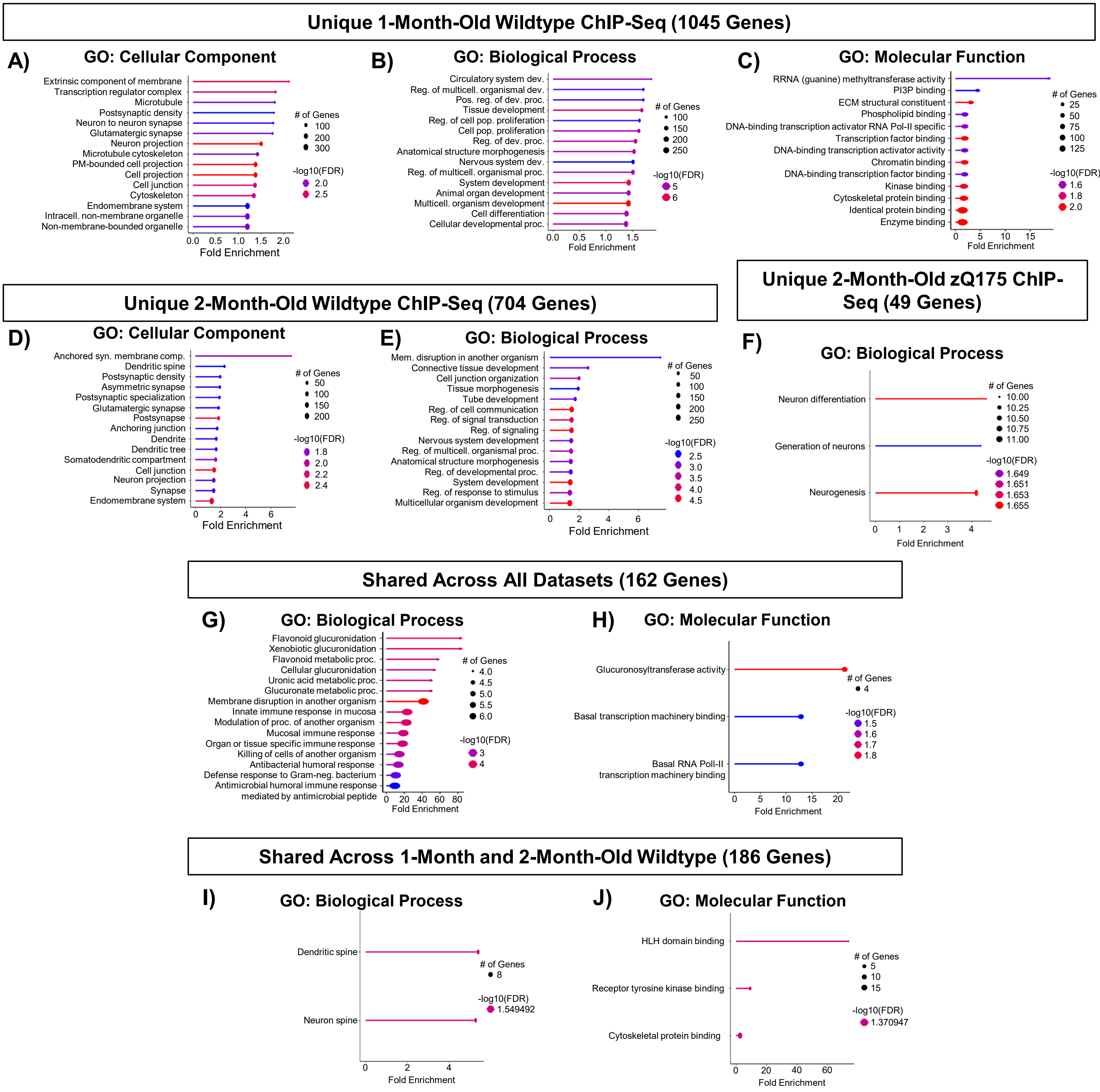


**Supplementary Figure 2**

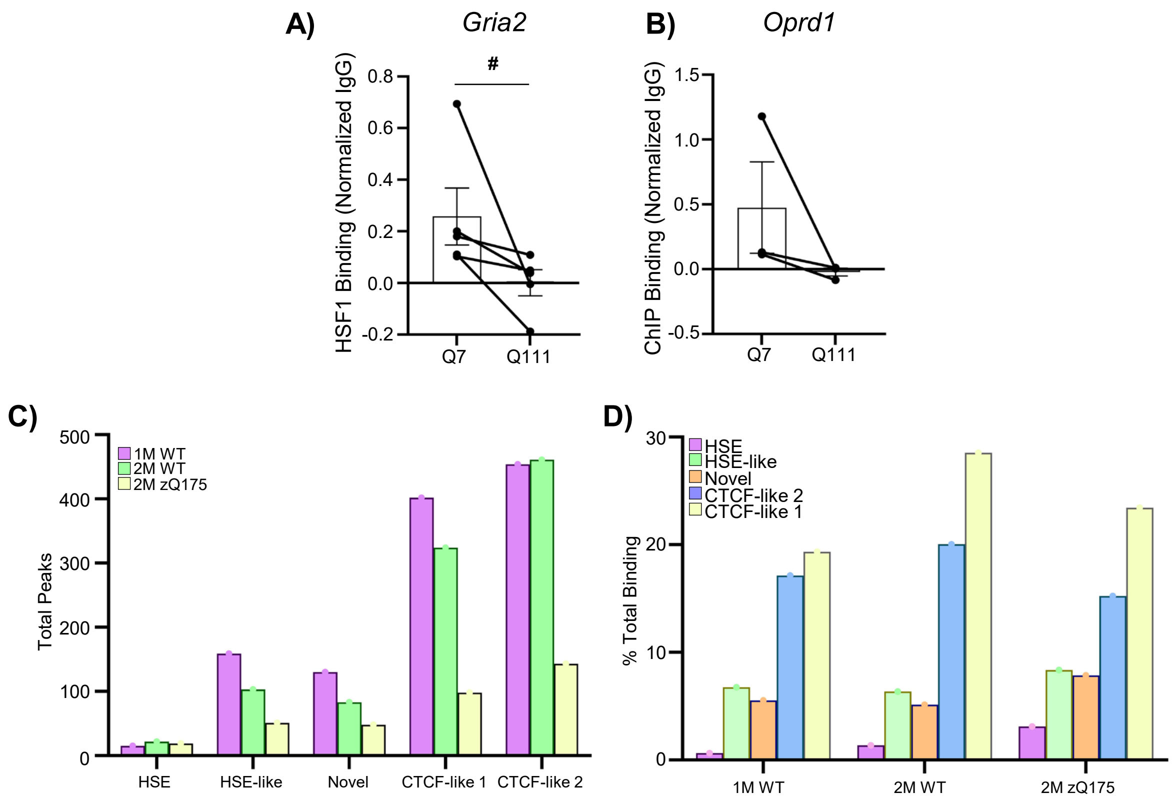


**Supplementary Figure 3**

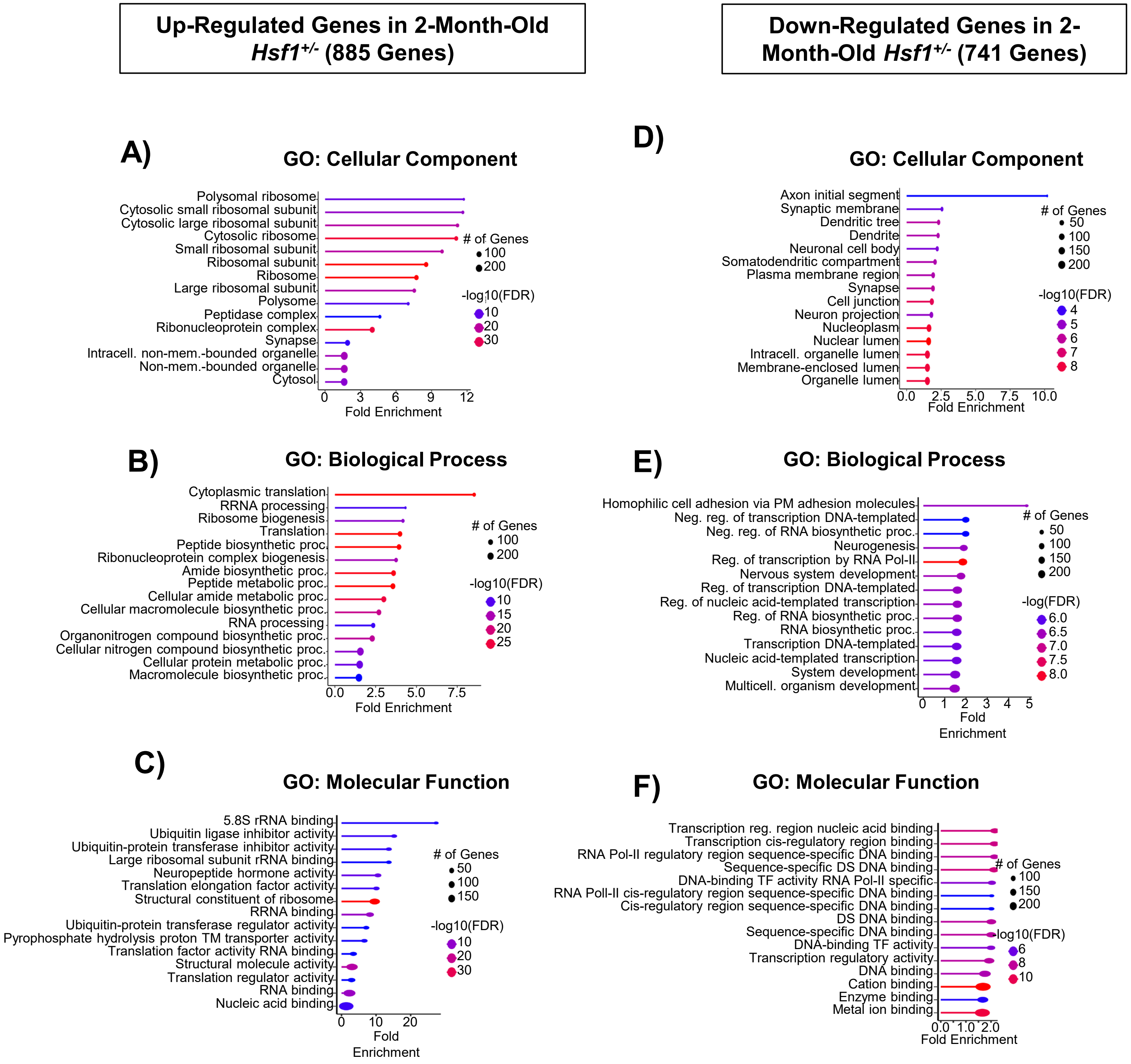


**Supplementary Figure 4**

#### A) Top Upstream Regulators of Differentially Expressed Genes in 2M HSF1 $^{+/-}$

| Upstream Regulator | Molecule Type | IPA Predicted Activation | Activation z-score | p-value | RNA | Protein |
|---|---|---|---|---|---|---|
| LARP1 | translation regulator | INHIBITED | -6.925 | 3.85E-39 | | |
| SPEN | transcription regulator | ACTIVATED | 6.083 | 5.52E-31 | | |
| FMR1 | translation regulator | INHIBITED | -3.786 | 8.90E-30 | * | |
| MLXIPL | transcription regulator | ACTIVATED | 6.066 | 8.38E-27 | | |
| YAP1 | transcription regulator | | 0.721 | 1.87E-17 | | |
| MYCN | transcription regulator | ACTIVATED | 4.29 | 1.58E-16 | | |
| CTNNB1 | transcription regulator | | -0.446 | 4.88E-14 | | |
| CREB1 | transcription regulator | | 1.462 | 1.64E-12 | | |
| MYC | transcription regulator | ACTIVATED | 5.695 | 4.30E-12 | | |
| HTT | transcription regulator | | 1.647 | 4.34E-12 | * | |
| RRP1B | transcription regulator | | | 1.26E-08 | | |
| IGF2BP1 | translation regulator | | -1.897 | 2.52E-08 | | |
| NFE2L2 | transcription regulator | ACTIVATED | 3.192 | 1.07E-07 | | |
| MAP3K12 | kinase | | | 1.21E-06 | | |
| ATN1 | transcription regulator | | | 7.36E-06 | | |
| EIF4E | translation regulator | | 2.119 | 3.27E-06 | * | |
| FUS | transcription regulator | | 0.555 | 4.22E-05 | | |
| FEV | transcription regulator | | 1.291 | 4.26E-05 | | |
| EIF6 | translation regulator | INHIBITED | -2.668 | 7.28E-05 | | |
| BMAL1 | transcription regulator | INHIBITED | -2.087 | 7.29E-05 | | |
| SALL4 | transcription regulator | | -0.146 | 1.18E-04 | | |
| HNF4A | transcription regulator | | 0.493 | 1.57E-04 | | |
| NFE2L3 | transcription regulator | | 1.444 | 1.68E-04 | | |
| MOS | kinase | | | 1.68E-04 | * | |
| NRF1 | transcription regulator | | 1.841 | 1.75E-04 | | |
| REST | transcription regulator | INHIBITED | -2.389 | 1.94E-04 | | |
| EOMES | transcription regulator | | -1.126 | 3.28E-04 | | |
| FUBP1 | transcription regulator | | 1.96 | 3.53E-04 | | |
| TP53 | transcription regulator | | 0.15 | 3.94E-04 | | |
| ERG | transcription regulator | | -0.499 | 4.07E-04 | | |

## B)

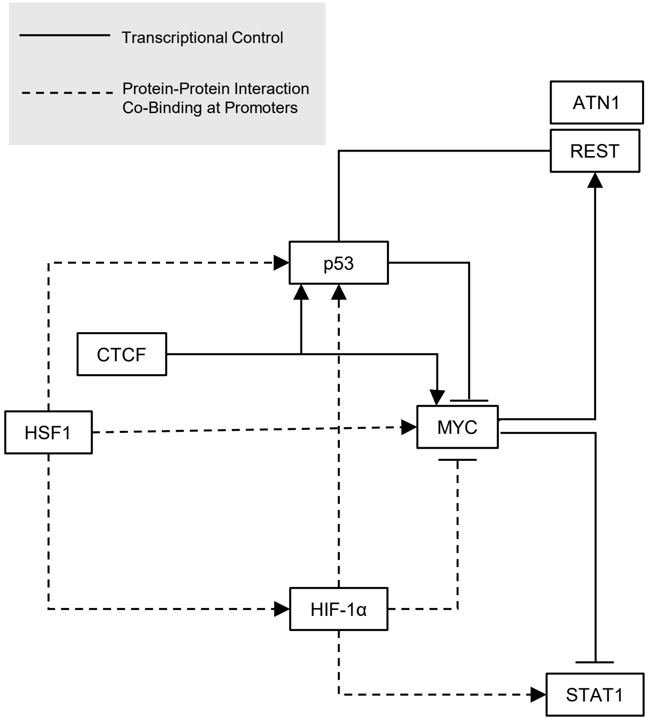


## C)

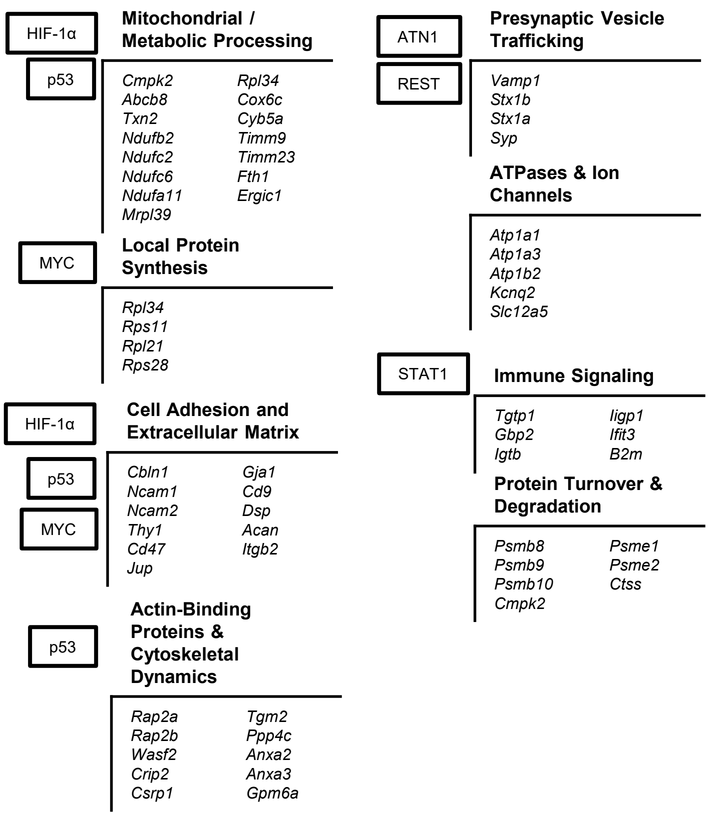


#### D) Top Upstream Regulators of Differentially Expressed Proteins in HSF1-cKO

| Upstream Regulator | Molecule Type | IPA Predicted Activation | Activation z-score | p-value | RNA | Protein |
|---|---|---|---|---|---|---|
| HTT | transcription regulator | | -1.181 | 5.15E-18 | * | |
| TYK2 | kinase | ACTIVATED | 3.138 | 2.22E-13 | * | |
| ATN1 | transcription regulator | | | 2.73E-10 | * | |
| TRIM24 | transcription regulator | INHIBITED | -2.69 | 1.27E-08 | | |
| NRF1 | transcription regulator | | -0.943 | 9.36E-08 | | |
| MYC | transcription regulator | INHIBITED | -2.085 | 3.38E-07 | | |
| PPP1R13L | transcription regulator | | 0.277 | 4.24E-07 | * | |
| SIRT1 | transcription regulator | INHIBITED | -3.055 | 5.71E-07 | | |
| MTOR | kinase | | -0.732 | 1.02E-06 | * | |
| HIF1A | transcription regulator | ACTIVATED | 2.032 | 1.25E-06 | | |
| BHLHE40 | transcription regulator | ACTIVATED | 2.496 | 1.27E-06 | * | |
| FMR1 | translation regulator | | -0.666 | 1.32E-06 | | |
| CTNNB1 | transcription regulator | | 1.46 | 1.34E-06 | | |
| PIK3CG | kinase | INHIBITED | -2.804 | 1.39E-06 | * | |
| INSR | kinase | | 1.739 | 1.49E-06 | * | |
| IGBP1 | phosphotase | | -1.732 | 2.15E-06 | | |
| TP53 | transcription regulator | | -0.225 | 3.05E-06 | | |
| STAT1 | transcription regulator | ACTIVATED | 3.482 | 5.88E-06 | | * |
| ARID1A | transcription regulator | | 1.321 | 6.72E-06 | * | |
| IRF7 | transcription regulator | ACTIVATED | 2.919 | 1.02E-05 | | |
| IRF1 | transcription regulator | ACTIVATED | 2.762 | 1.08E-05 | | |
| STAT3 | transcription regulator | | 0.303 | 1.97E-05 | | |
| FUS | transcription regulator | | | 2.32E-05 | | |
| TCFL2 | transcription regulator | | 0.943 | 2.64E-05 | | |
| CITED2 | transcription regulator | INHIBITED | -2.203 | 3.87E-05 | | |
| ATM | kinase | | 0.33 | 4.12E-05 | * | |
| TEAD1 | transcription regulator | | 0.113 | 4.14E-05 | | |
| BCKDK | kinase | | | 4.17E-05 | | |
| REST | transcription regulator | INHIBITED | -2.207 | 4.85E-05 | * | |
| TARDBP | transcription regulator | | 0.128 | 5.45E-05 | | |

2 1 0 -1 -2

**Supplementary Figure 5**

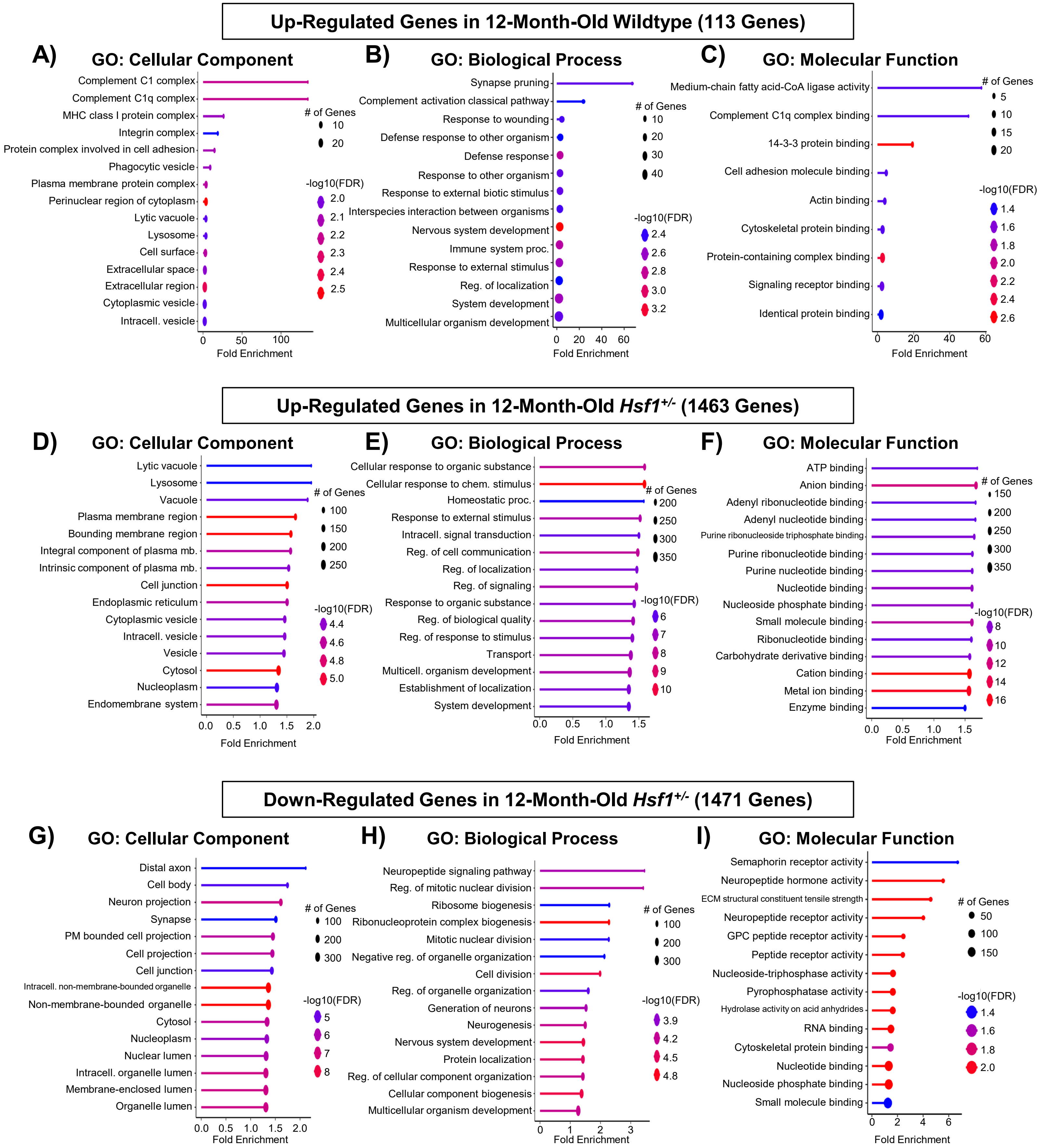


**Supplementary Figure 6**

#### Down-Regulated Proteins HSF1-cKO Synaptosomes ShinyGO Database (46 Proteins)

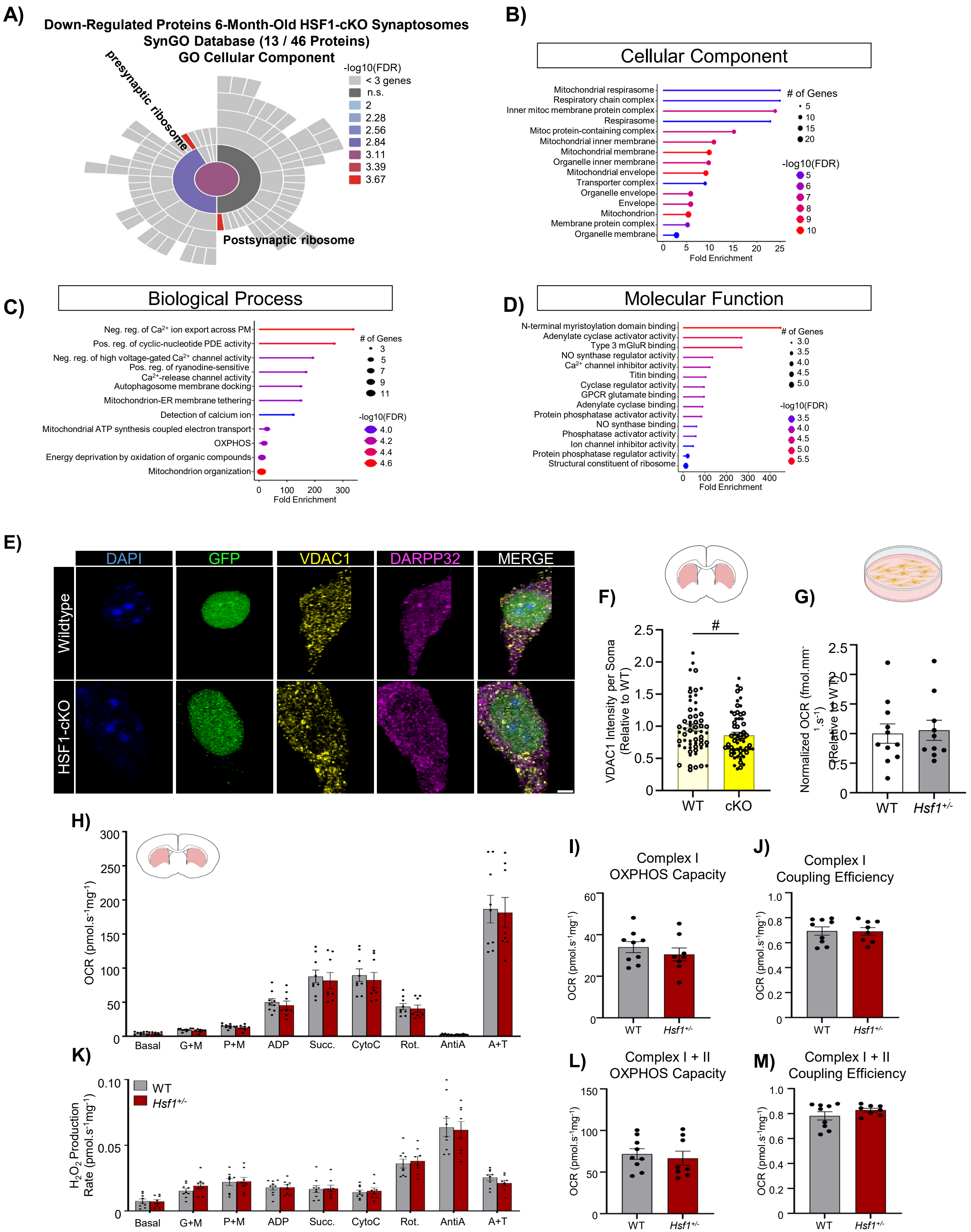


#### Supplementary Figure 7

### Up-Regulated Proteins HSF1-cKO Synaptosomes ShinyGO Database (156 Proteins)

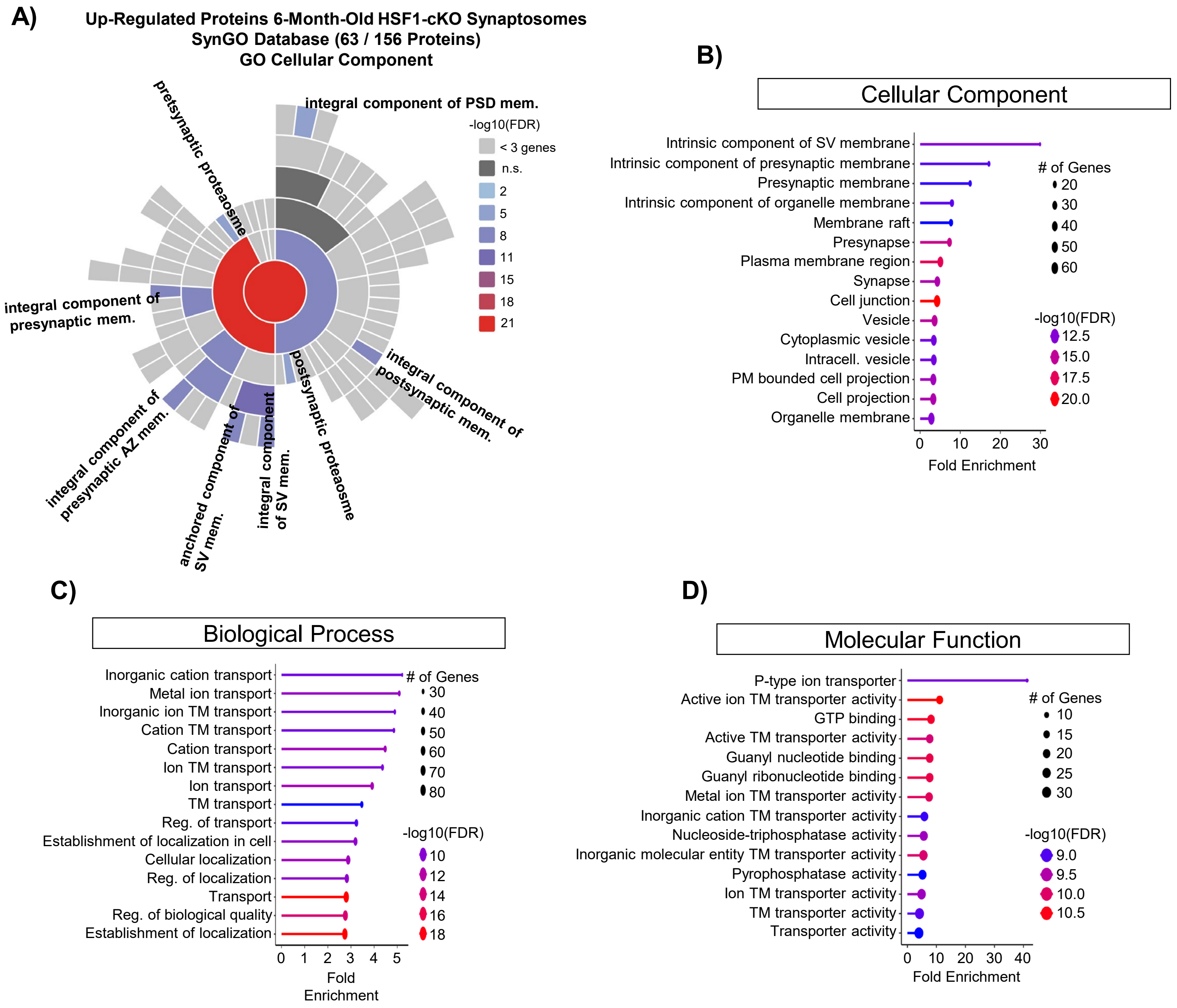


**Supplementary Figure 8**

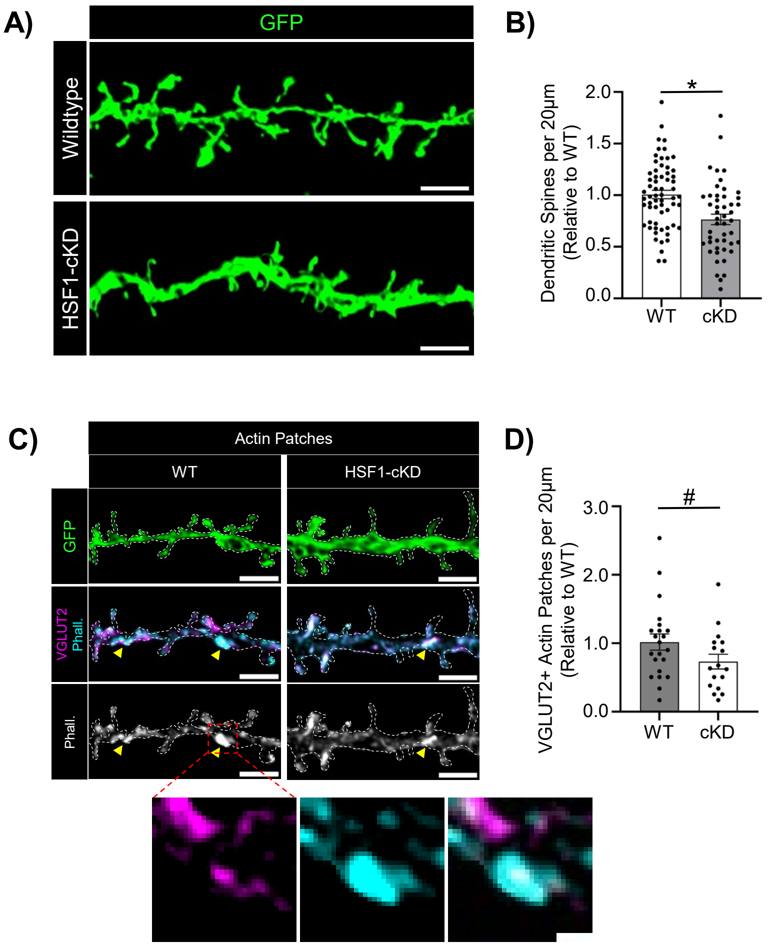


**Supplementary Figure 9**

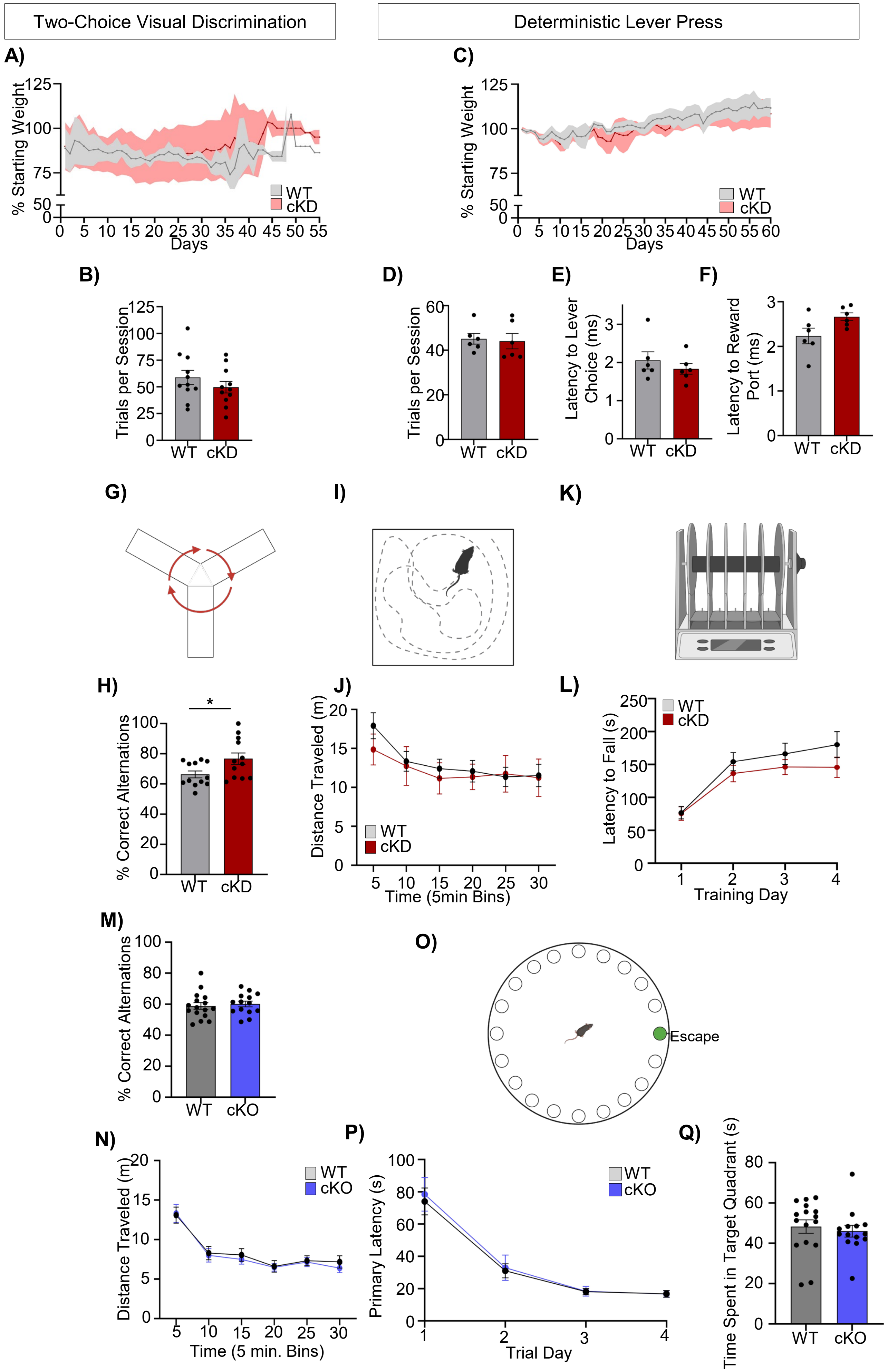


**Supplementary Figure 10**
